# A new method for a priori practical identifiability

**DOI:** 10.1101/2022.10.20.511900

**Authors:** Peter Thompson, Benjamin Jan Andersson, Nicolas Sundqvist, Gunnar Cedersund

## Abstract

**Background and objective:** Practical identifiability analysis – to determine whether a model property can be determined from given data – is central to model-based data analysis in biomedicine. The main approaches used today all require that the coverage of the parameter space be exhaustive, which is usually not possible. An attractive alternative could be to use structural identifiability methods, since they do not need such a parameter coverage. However, current structural methods are unsuited for practical identifiability analysis, since they assume that all higher-order derivatives of the measured variables are available, and we do not know the implications of this assumption. Herein, we provide new definitions and methods that allow for this assumption to be relaxed.

**Methods:** The new methods and definitions are valid for ordinary differential equations, and use a combination of differential algebra and modulus calculus, implemented in Maple.

**Results:** We introduce the concept of (ν_1_, …, ν_*m*_)-identifiability, which differs from previous definitions in that it assumes that only the first ν_*i*_ derivatives of the measurement signal *y*_*i*_ are available. This new type of identifiability can be determined using our new algorithm, as is demonstrated by applications to various published biomedical models. Our methods allow for identifiability of not only parameters, but of any model property, i.e. observability. These new results provide further strengthening of conclusions made in previous analysis of these models. Importantly, our analysis can for the first time quantify the impact of the assumption that all derivatives are available in specific examples. If one e.g. assumes that only up to third order derivatives, instead of all derivatives, are available, the number of identifiable parameters drops from 17 to 1 for a Drosophila model, and from 21 to 6 for an NF-κB model. In both these models, the previously obtained identifiability is present only if at least 20 derivatives of all measurement signals are available.

**Conclusions:** Our results demonstrate that the assumption regarding availability of derivatives done in traditional structural identifiability analysis causes a big overestimation regarding the number of parameters that can be estimated. Our new methods and algorithms allow for this assumption to be relaxed, which moves structural identifiability methodology one step closer to practical identifiability analysis.

## 1 Background

Biology and medicine involve the study of a myriad of time-dependent variables, which cross-talk with each other in complex networks. To deal with this complexity, one often makes use of mechanistic mathematical models, often formulated using ordinary differential equations (ODEs). These ODEs represent hypotheses regarding the biological mechanisms that can explain given biological data [1]. The model-based data analysis of such real data revolves around a correct uncertainty analysis of parameters and predictions: to what degree can one determine the parameters and predictions of interest? This uncertainty analysis is formally called identifiability analysis, if it deals with parameter uncertainty, and observability analysis, if it deals with other model properties, such as states and composite variables [2]. Both observability and identifiability analysis can be done in many different ways, but they are generally subdivided into two types of approaches: structural and practical.

Practical identifiability analysis revolves around the specific data one has collected in a specific situation ([2]). In other words, practical identifiability analysis looks at the real situation, where the data has noise, and where the experiment might not have been done in an optimal way to generate information about all parameters. Historically, one has often used sensitivity-based methods to study practical identifiability, such as analysis of the Fisher Information Matrix ([3]), and the covariance of the parameters ([4]). In the last 10 years, these methods have been increasingly replaced by Markov Chain Monte Carlo (MCMC) and profile-likelihood based methods ([5], [2], [4]). The benefit of these more modern methods is that they study global properties, and do not have to assume that the system is identifiable to assess identifiability ([3], [2]). These methods are highly useful and flexible, but they still have limitations. Most importantly, they rely on the exhaustive coverage of the entire parameter space, either using the MCMC sampling or using the optimization step in profile likelihood. For large models, this optimization step is difficult and is time-consuming even for an expert in the field ([6], [7]). This difficulty will always leave the analysis results with an inherent doubt: if the coverage of the parameter space fails, the degree of identifiability (the calculated confidence intervals) is not correct.

Structural identifiability analysis has historically been the mirror image of practical identifiability analysis. Structural methods normally do not take a specific data set or the noise level into account. Instead structural identifiability analysis only considers the form of the equations, including the measurement equations. Furthermore, structural identifiability analysis does not rely on an exhaustive coverage of the entire parameter space. Instead, structural identifiability analysis makes use of powerful theories from differential algebra and differential geometry to investigate whether there are structural limits for whether a parameter or a model property ever can be determined from a given measurement. Methods have been developed to determine which states and parameters are locally identifiable ([8]) and globally identifiable ([9], [10]), to discover parameter combinations that are identifiable ([11], [6], [12], [13], [14]), and to suggest modifications in the modeling process based on identifiability ([15], [16]). In the last two decades, powerful methods have been developed that can do such an analysis also for relatively large biological models, featuring 50 to 100 states and parameters ([16]).

Because structural identifiability now can be applied to real, large models, it would be highly interesting if they also could perform practical identifiability analysis, where the limitations of a specific dataset are taken into account. The probably most prominent such assumption is that current methods assume that all derivatives of all measurement signals can be estimated from the data. In practical situations, with real data, this assumption is never fulfilled. It is therefore a critical flaw that we are lacking definitions and methods to analyze the consequences of this unfulfilled assumption.

In summary, there is therefore a need to develop new methods that combine the strengths of the two approaches: A method that does not rely on too idealistic assumptions regarding the real situations, but that also does not rely on optimization and coverage of the entire parameter space. However, such methods have not yet been proposed. Herein, we introduce a new type of practical *and* structural identifiability, which puts an upper limit on how many times each measurement signal can be differentiated. We also introduce a new algorithm that can calculate this new type of identifiability, and apply this to a series of published models. For all studied examples, traditional structural identifiability methods have widely overestimated the number of identifiable parameters: with a maximum of 3 derivatives available, more than 90% of the previously identifiable parameters lose their identifiability.

## 2 Results and Discussion

### 2.1 Problem: there is an upper limit for how many derivatives can be estimated from biological data

There is usually an upper limit for the number of derivatives of an output signal that can be estimated from data. This is illustrated in the following simulation. Consider the system

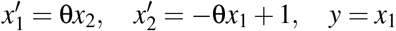

where *x*_1_ and *x*_2_ are state variables, θ is a constant parameter, and *y* is an observable output variable; that is *y* is measurable by experiments but *x*_1_, *x*_2_ and θ are unknown. Suppose that, unbeknownst to the observer, θ = 1, *x*_1_(0) = 10, and *x*_2_(0) = 0. Consequently the true signal of *y* is given by *y*(*t*) = 1 + 9 cos(*t*). However, in practice, the measured signal will not show a smooth curve but rather a discrete time series that also has added noise. An illustration of this measurement signal can be found in Figure 1A, where 10 realizations of the function *y*(*t*) + *N*(0, σ = 2), where the second term represents normally distributed noise with mean 0 and standard deviation 2, are depicted alongside the true function *y*(*t*) (Figure 1A). The number of derivatives that can accurately be estimated can be evaluated by deriving the function of a polynomial interpolation of these data points and comparing these estimated derivatives to the derivatives of the true function (Figure 1B). More specifically, this example shows a 6^*th*^ degree polynomial interpolation for t-values ranging from 0 to 6 in increments of 0.5. Subsequently, the 0^*th*^ to the 5^*th*^ derivatives of these interpolations are then compared to the derivatives of the true function *y*(*t*). The number of derivatives that can accurately be estimated is dependent on the magnitude of noise in the system. More specifically, the magnitude of the added noise increases the number of accurately estimated derivatives declines (Figure 1C). To illustrate this dependency, the example above was repeated with the added noise sigma ranging from 0.1 to 10 in increments of 0.1 (σ = [0.1, 0.2,…, 10]) and with 1000 data realizations generated at each noise level. Figure 1C shows the average number of derivatives that could accurately be estimated at each noise level. A derivative is considered accurate if the square root of the sum of squared residuals does not exceed the amplitude of *y*. From this figure, it is clear that the number of derivatives that can accurately be estimated diminishes rapidly as the signal-to-noise ratio approaches 1.

**Figure 1:**
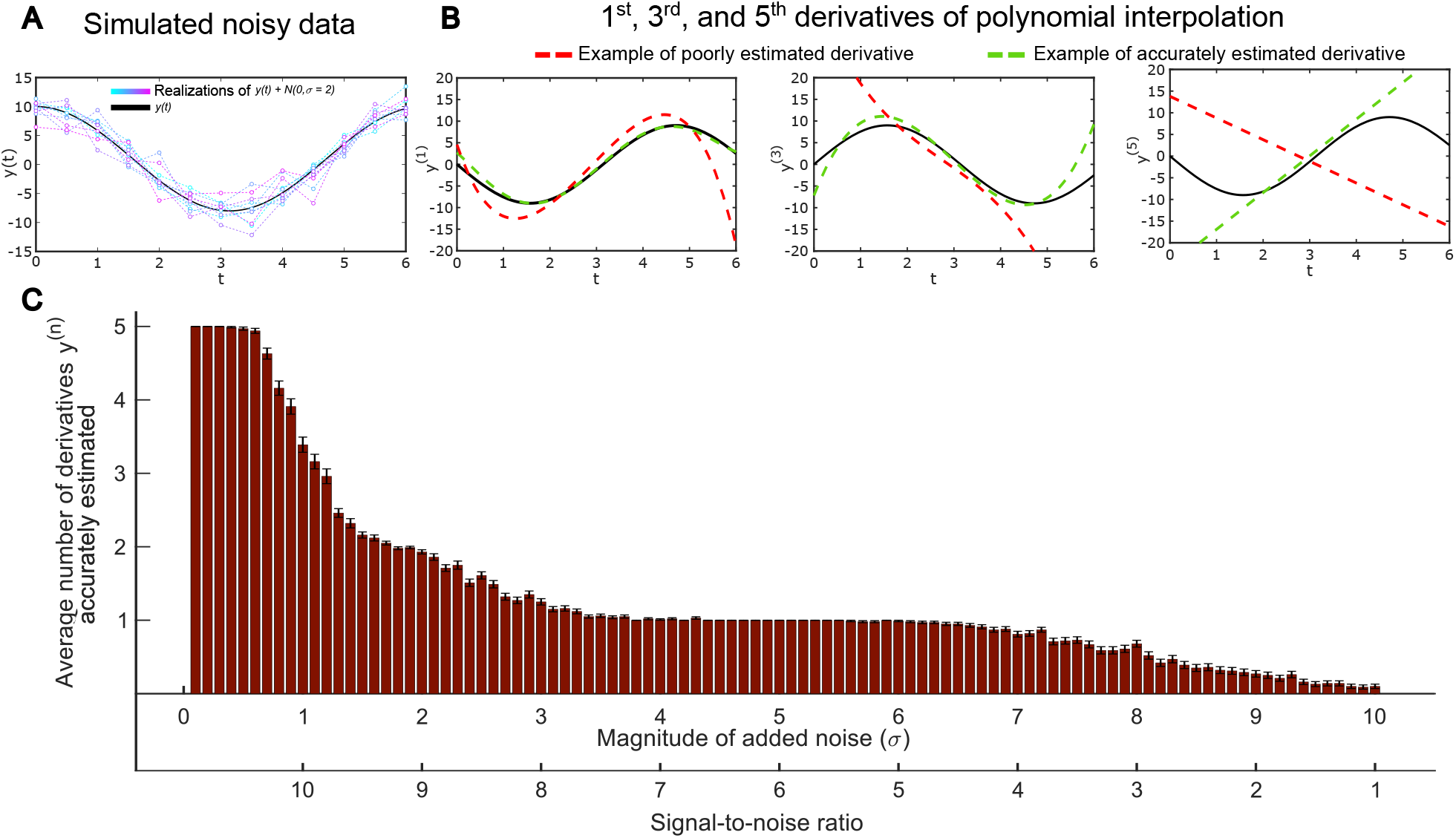
Illustration of the upper limit for how many derivatives can be estimated from noisy data. **A:** shows examples of 10 different realizations of noisy data (colored lines) and the true function *y*(*t*) = 1 + 9 cos(*t*) (back line) **B:** Illustrations of the 1^*st*^, 3^*rd*^, and 5^*th*^ derivatives of the function *y*(*t*) with examples of a poorly estimated derivative (red line) and a accurately estimated derivative (green line). **C:** The relationship between the average number of accurately estimated derivatives and the magnitude of the added noise σ/the signal-to-noise ratio. The bars show the mean number of accurately estimated derivatives at each noise level and the error bars show the standard error of the mean (SEM).

The following example illustrates how this lack of availability of all derivatives of *y* is associated with an unrealistic assessment of whether parameters are identifiable.

#### Example 2.1.

Consider the system

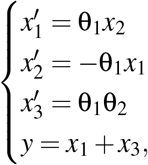

where *x*_1_, *x*_2_, and *x*_3_ are (unobserved) state variables, θ_1_ and θ_2_ are constant parameters, and *y* is an (observed) output variable.

Running the algorithm from [8] shows that *x*_1_, *x*_2_, *x*_3_, θ_1_, and θ_2_ are locally identifiable. Let us attempt to estimate θ_1_ from observation of *y*. A relation among θ_1_ and *y* over ℂ is 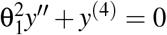.Thus we can estimate θ_1_ by using our observed values of *y*^*tt*^ and *y*^(4)^ at some time *t*_0_:

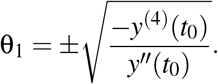

In practice, the measured signal *y* may be too noisy to give an accurate estimation of *y*^(4)^. In such a situation, θ_1_ is not identifiable from the given data, even though a structural analysis says that it is. Suppose now that we know that our measurements are good enough to give reliable estimates of *y*″ but of no higher derivative of *y*. To use structural methods to determine the degree of identifiability in such a situation, we need new definitions and methods.

#### Remark 2.2.

Our approach assumes measurements of several derivatives of the output signal are made at a single time. In practice it is more common to measure the 0-th derivative of the output signal at several times. Having measurements with some uncertainty of *y*(*t*_0_), …, *y*(*t*_*N*_) is equivalent to having measurements with some uncertainty of *y*(*t*_0_), …, *y*^(*N*)^(*t*_0_). Hence this aspect of our approach does not limit its generality.

For Example 2.1, one can estimate θ_1_ using the relation 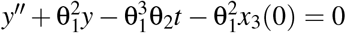 together with measurements of *y* and *y*″ at three different times. However knowledge of *y, y*′, *y*″ at three times can be used to estimate *y, y*′, *y*″, *y*^(3)^, *y*^(4)^ at a single time.

### 2.2 A new definition for practical and structural identifiability: (ν_1_, …, ν_*m*_)*-identifiable*

We now provide an intuitively straightforward definition that is useful for understanding the new methods. In Section 5, we provide a precise definition, together with analytical proofs of all stated method properties.

Let *ℓ, n, m*, and *r* be positive integers. Let **θ** = (θ_1_, …, θ_*ℓ*_), **x** = (*x*_1_, …, *x*_*n*_), **y** = (*y*_1_, …, *y*_*m*_), and **u** = (*u*_1_, …, *u*_*r*_). Let **f** = (*f*_1_, …, *f*_*n*_) and **g** = (*g*_1_, …, *g*_*m*_) be tuples of rational functions in **x, u**, and **θ** over ℂ. This setup determines a class of systems of ODEs with initial conditions, or a model class:

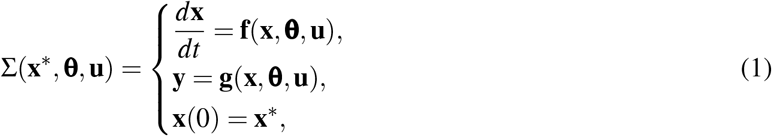

where **x** are state variables, which usually correspond to time-varying concentrations or physical properties; **x**^∗^ are the initial values of the state variables; **θ** are constant parameters, e.g. corresponding to rate constants or volumes; **u** are input variables, which are observed and typically controlled by the experimentalist; and **y** are output variables, which are observed experimentally. With these notations in place, the new definition is roughly given by:

#### Definition 2.3

(roughly stated). Let ν_1_, …, ν_*m*_ be non-negative integers. Rational expression *h*∈ ℂ(**x, θ**) is said to be (ν_1_, …, ν_*m*_)*-identifiable* if perfect knowledge of the first ν_*i*_ derivatives of *y*_*i*_ at a particular time allows us to determine the value of *h* at that point in time, up to a finite set.

This definition, which is more technically stated in Definition 5.4, provides the essence of the new definition: given only a limited set of derivatives of the different measurement signals, can we determine the parameter θ_*i*_ to be locally identifiable, i.e. as belonging to a finite set? This definition is analogous to traditional definitions of structural identifiability, with the only addition being that we now only have access to a finite set of derivatives. In other words, in traditional structural identifiability analysis, no such constraints on the number of derivatives available is present. This new definition is closer to practical identifiability since, with real biological data, one can only estimate a few derivatives from the measurement data.

### 2.3 Solution: our new algorithms for calculating practical structural identifiability

For most models, it would be difficult to prove the (ν_1_, …, ν_*m*_)-identifiability of a given parameter directly from the definition. We provide Algorithm 1, which can do this using only straightforward algebraic computations. The algorithm applies to any rational combination of states and parameters, of which an individual parameter is a special case.

#### Algorithm 1

Determine whether a given element of ℂ(**x, θ**) is **ν**-identifiable

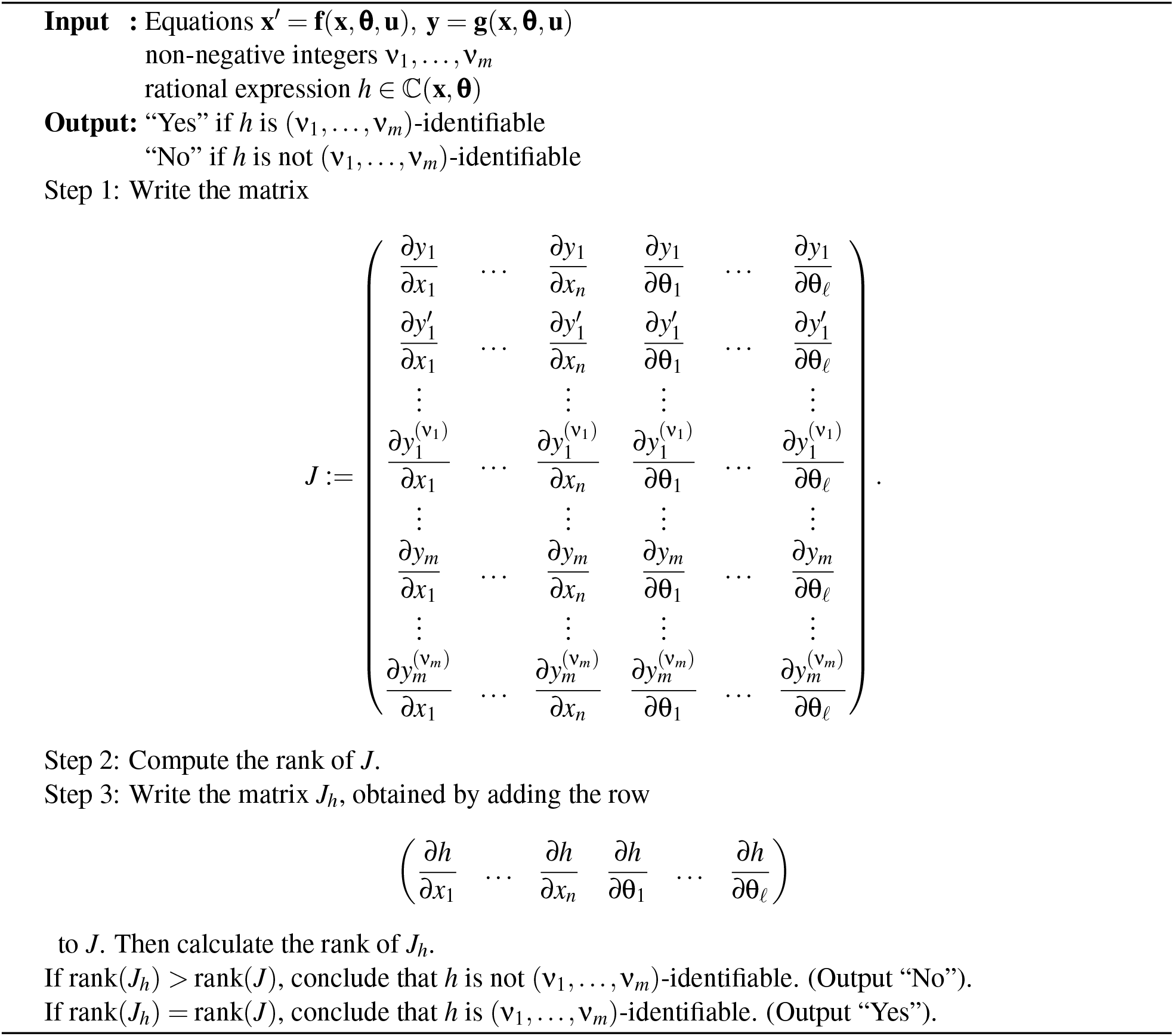

#### Algorithm 2

Determine whether a given element of **x** ∪ **θ** is **ν**-identifiable

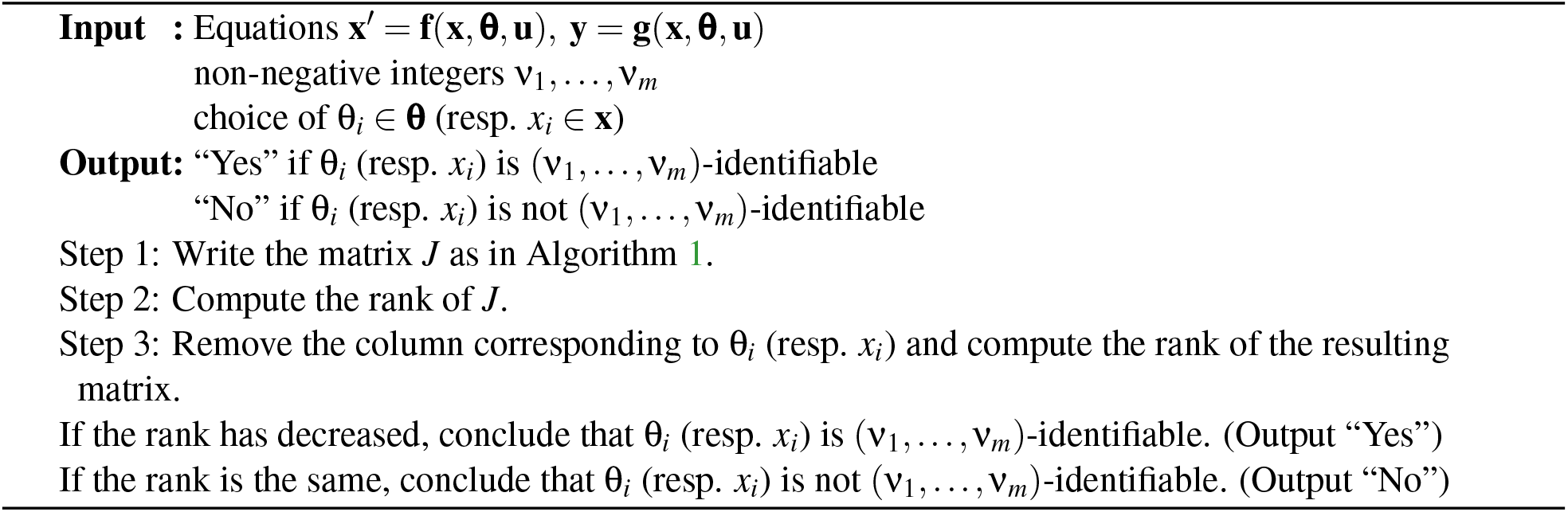

In the case where one is interested in just an individual parameter or state, one can perform Algorithm 2, where one column is removed instead of a row added, resulting in a smaller matrix.

The precise definition of **ν**-identifiability is given in Definition 5.4, and we prove that Algorithms 1 and 2 are correct in section 5.2.

The symbolic calculations can be impractically slow for all but the smallest models. Therefore we have also provided probabilistic analogs of these algorithms that are much faster in practice. These are presented and their correctness is proven in section 5.3.

### 2.4 Intuitive understanding of the new algorithm

Each row of *J* represents how an output, or one of its derivatives, varies with respect to small changes in each state and parameter. If the last row of *J*_*h*_ is a linear combination of the other rows, then the variation of *h* can be accounted for as a combination of the variations of the output derivatives. This, in turn, means that it seems like *h* can be expressed in terms of the known signals, i.e. that *h* is identifiable. Conversely, if the last row of *J*_*h*_ is linearly independent of the other rows, then the variation of *h* cannot be explained in terms of that of the available derivatives of the outputs, and one should not expect to be able to determine *h* from the available outputs and their derivatives.

### 2.5 Examples

#### 2.5.1 Algorithm 1 determines identifiability of rational quantities, which is important in hypothesis testing

Algorithm 1 (and its faster probabilistic version Algorithm 3) can be used to analyze so-called core predictions (see [2]). A core prediction is a well-determined property, i.e. a model prediction with a small uncertainty. Such core predictions are therefore often tested in future experiments, and are a central part of model-based hypothesis testing. Algorithm 1 can test whether or not such a model property really can be well-determined, given the available data. We illustrate this new possibility in the following two examples.

##### Example 2.4.

Figure 2 shows a model of insulin (ins) binding and activation of insulin receptor (IR). This model was originally presented in [17], and is explained more in detail therein and in the supplementary materials. This model was one of several hypotheses tested in the article. To draw one of the conclusions in that article, the authors looked at the proportion of insulin receptor bound to an internal membrane:

**Figure 2:**
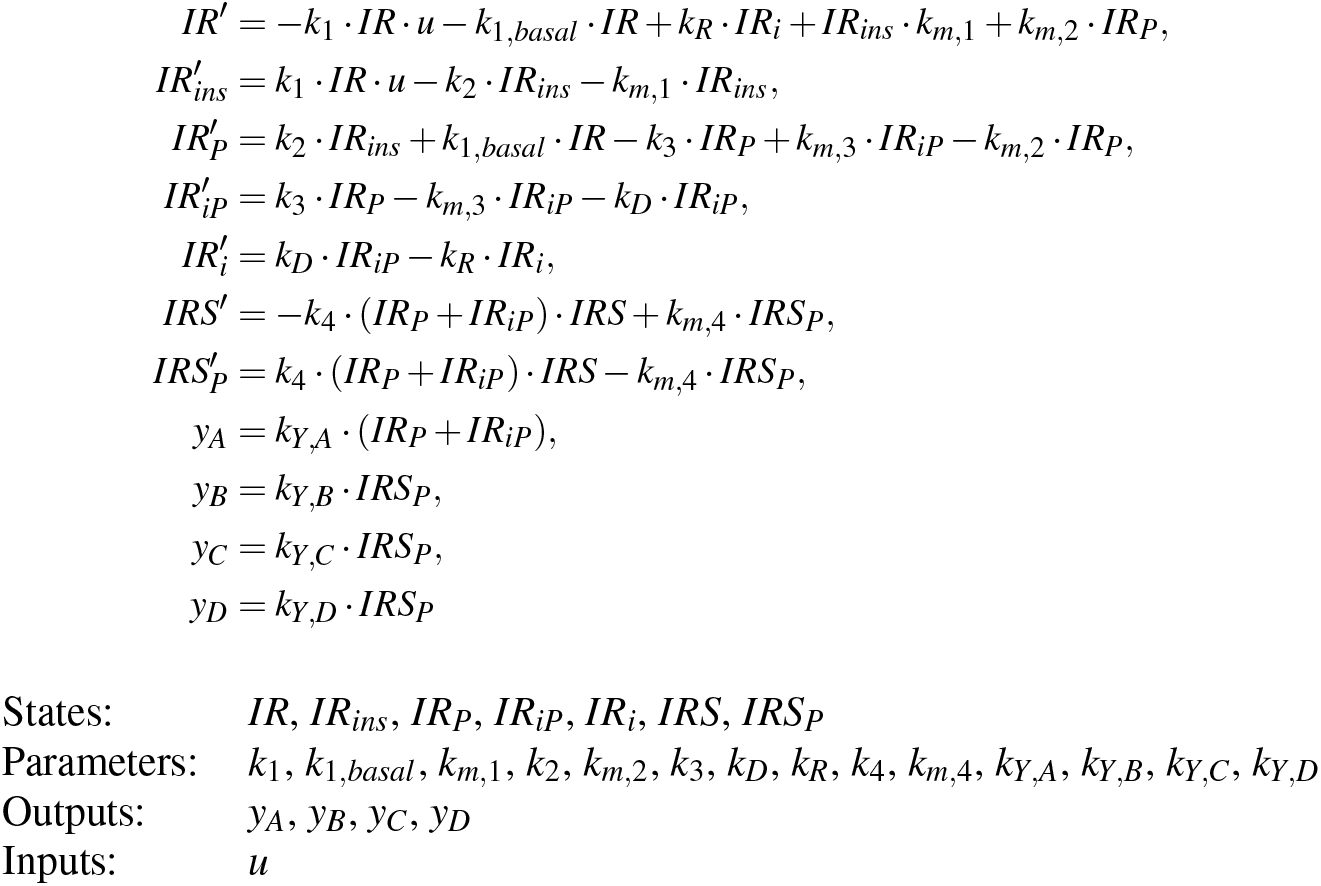
Model for insulin uptake. (For more details, see “Supplementary Materials”.)

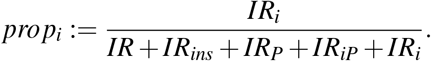

It was predicted that *prop*_*i*_ should be between .55 and .80. Despite the fact that there are four outputs, none of the seven states (insulin receptor or substrate quantities at different locations) are identifiable, so it is not immediately obvious whether *prop*_*i*_ can be estimated from the measurements. Algorithm 3 with ν_*i*_ = *n* + ℓ−1 for all *i* shows that this quantity is indeed single-experiment locally identifiable (see Definition 5.8 and Corollary 5.10). The authors were able to experimentally determine the proportion to be 0.023 ± 0.008. Thus they were able to reject the model, which was the conclusion in that step in the paper.

In summary, our analysis shows that the proportion of receptors that are bound to an internal membrane indeed is an identifiable model property. This new result holds from a structural point of view, without assuming that all derivatives are known. These results also show this without relying on optimization or exhaustive parameter sampling, as is done using the traditional practical identifiability analysis done in the original papers [17] and [2]. This new result thus strengthens and confirms the conclusion drawn in the original paper.

##### Example 2.5.

Figure 3 shows the model of the interaction between liver and pancreatic cells from [18]. There are seven states, *x*_1_, …, *x*_7_, two outputs, *y*_1_, *y*_2_, and no inputs. Numerical values were inserted for parameters whose values were known from operating specifications or estimated from the literature. The remaining parameters, here denoted by θ_1_, …, θ_9_, were to be determined from experimental data. One quantity of interest was

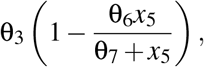

representing hepatic insulin sensitivity (see [18, Figure 4 Plot E]). Algorithm 3 tells us that this quantity is indeed identifiable. Moreover, by using different values of (ν_1_, ν_2_) we find that it is (6, 5)-identifiable.

**Figure 3:**
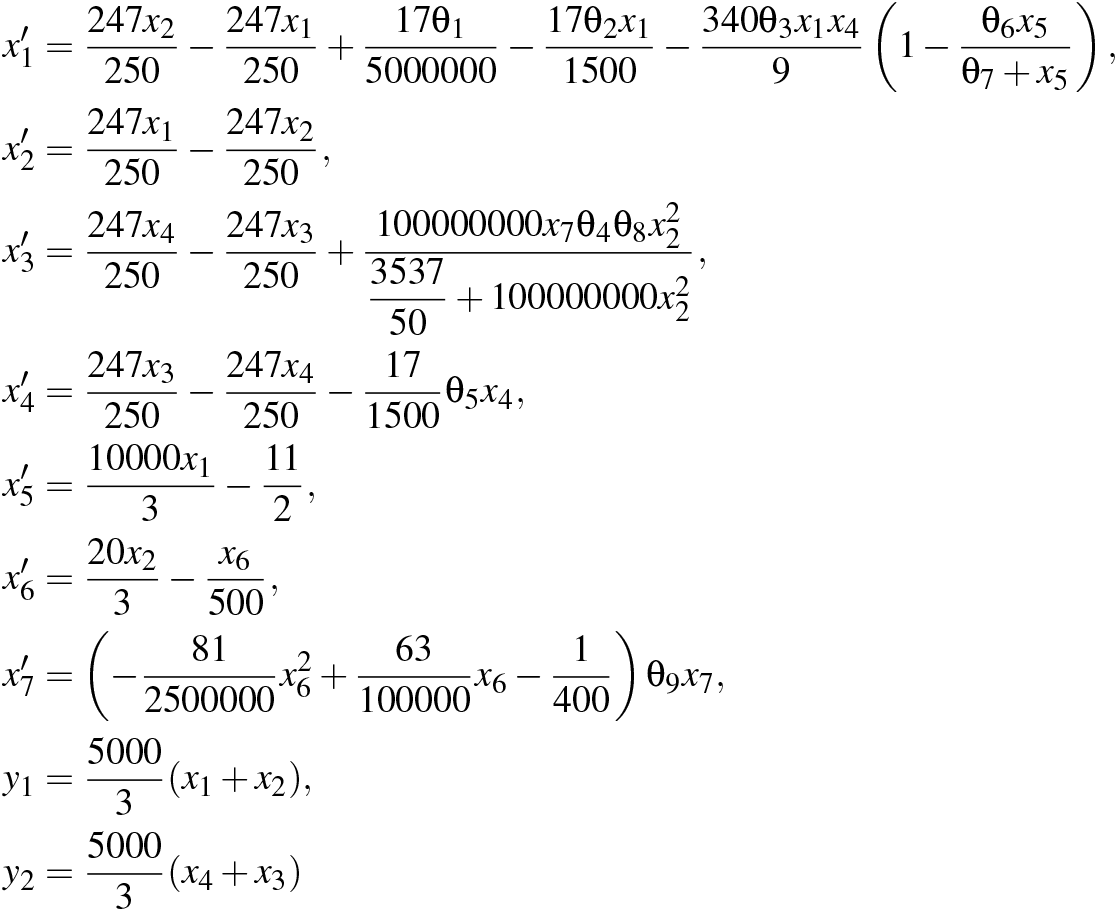
Model for interaction between liver and pancreatic cells. (For more details, see “Supplementary Materials”.)

**Figure 4:**
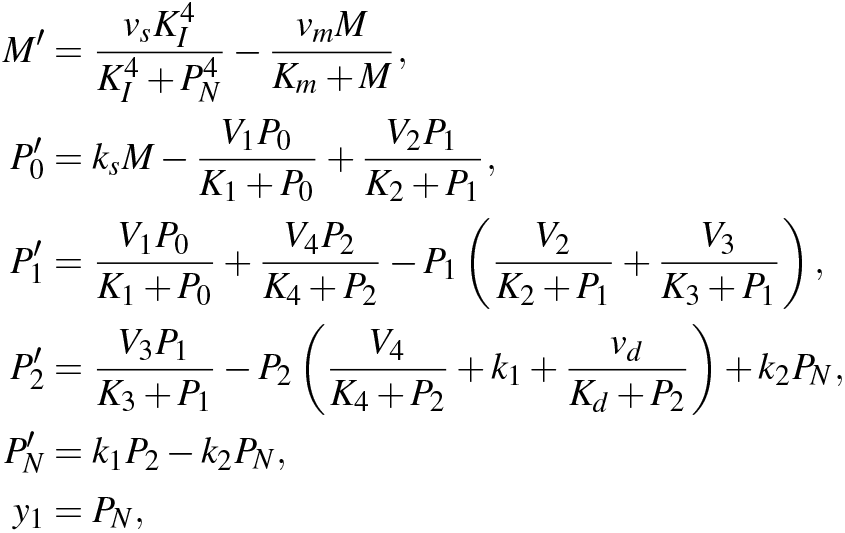
Model for Drosophilia period protein. (For more details, see “Supplementary Materials”.)

#### 2.5.2 Identifiability tends to increase with increased order of observability

It is typically the case that as we increase the number of derivatives that can be reliably estimated, the number of identifiable parameters increases, often dramatically. Therefore assuming that all derivatives are available can give a large overestimate of the number of identifiable parameters. How large this overestimate is can now be determined by our new algorithms. We illustrate this capability in the following two examples.

##### Example 2.6

(Drosophila period protein). Figure 4 gives the model that was used in [19] to model Drosophila period protein. A local structural identifiability analysis done in [8, p. 737] shows that all states and parameters other than *M, v*_*s*_, *v*_*m*_, *K*_*m*_, and *k*_*s*_ are identifiable. However, Algorithm 4 shows that only *P*_*N*_ is identifiable using 19 or fewer derivatives of *y*_1_. If in addition we observe *M*, it was shown in [8] that all states and parameters are identifiable. However Algorithm 4 shows this is the case only if at least 16 derivatives of the outputs are available. Figure 5 shows the number of states and parameters that are identifiable as a function of the maximum number of derivatives of each output that are available; for example ν_*max*_ = 3 means that exactly 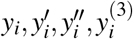 are available for all *i*. Blue circles represent output set {*y*_1_ = *P*_*N*_} and red squares represent output set {*y*_1_ = *P*_*N*_, *y*_2_ = *M*}.

**Figure 5:**
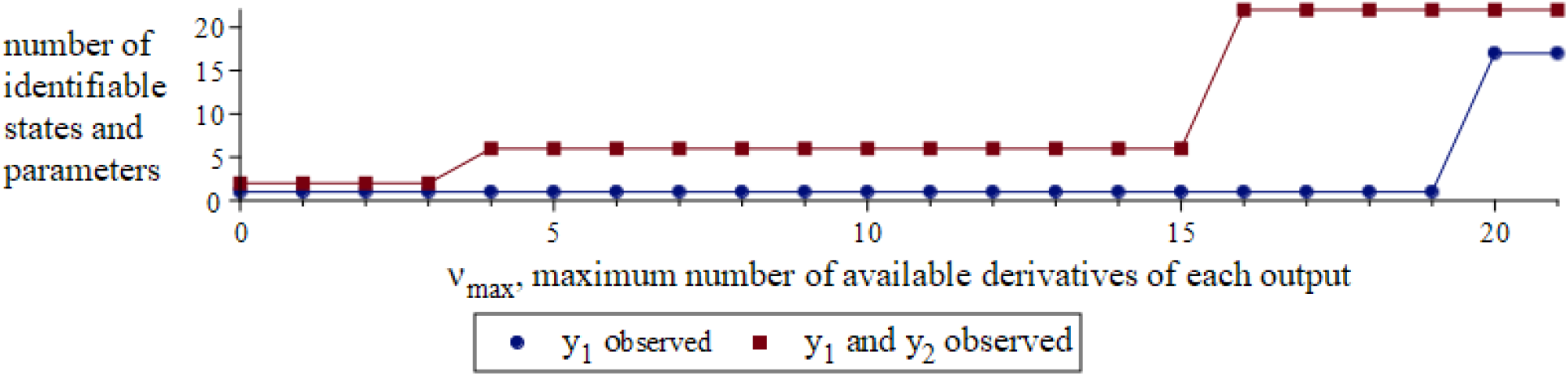
Variation of the number of identifiable parameters with maximum number of available output derivatives in a model of Drosophila period protein. The horizontal axis shows 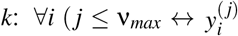 can be reliably estimated). The vertical axis shows the number of identifiable states and parameters in the model. Blue circles represent the case with only one output: *y*_1_ = *P*_*N*_. Red squares represent the case with two outputs: *y*_1_ = *P*_*N*_, *y*_2_ = *M*.

##### Example 2.7

(NF-κB). We consider a model of NF-κB regulatory module, as presented in Figure 6 below. This model was first introduced in [20], and an identifiability analysis was done in [21]. The model has 15 states, 28 parameters, 4 outputs, and 0 inputs.

**Figure 6:**
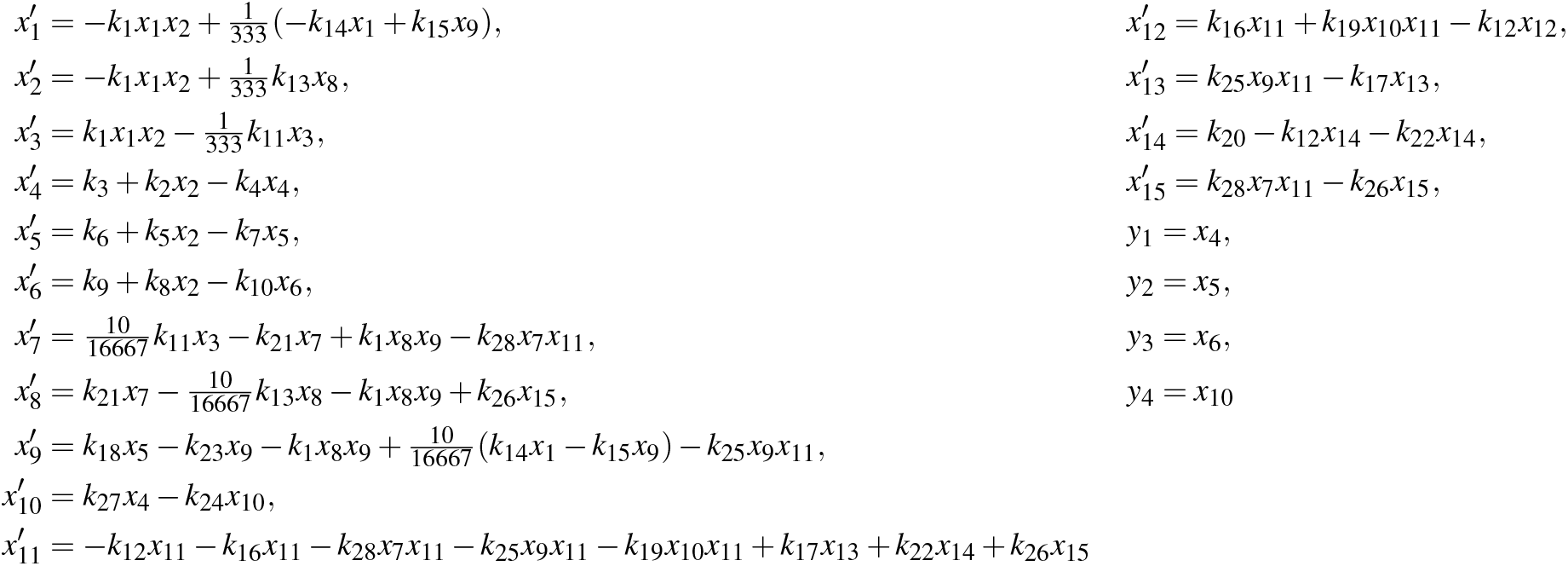
Model for NF-κB regulatory module. (For more details, see “Supplementary Materials”.)

The analysis in [21] shows that 21 states and parameters are identifiable, assuming that all derivatives of the four outputs can be estimated. However, using Algorithm 4, we found that if no more than 26 derivatives are available, then only 9 states and parameters are identifiable. Further analysis is shown in Figure 7.

**Figure 7:**
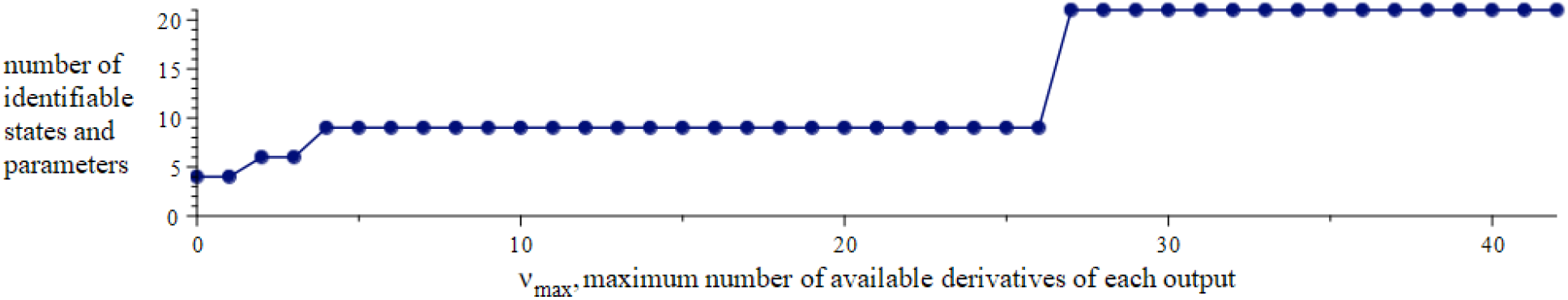
Variation of the number of identifiable parameters with maximum number of available output derivatives in a model of NF-κB regulatory module. The horizontal axis shows 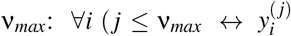 can be reliably estimated). The vertical axis shows the number of identifiable states and parameters in the model.

### 2.6 Relation to other available structural approaches to determine practical identifiability

We present a new method for determining structural identifiability in a more realistic setting, where only a given finite number of derivatives are available for each measurement signal. While this is novel, there are a few related methods that should be mentioned.

Most structural identifiability methods assume that all derivatives of outputs are measurable and all derivatives of all outputs are sufficiently varying. The latter condition is not always satisfied in practical modeling as, for example, a constant function may be used as an input, and this may lead to incorrect implementation of a model. In [22] a method for determining how many derivatives of the inputs must be non-zero was given, and an implementation was presented in [23].

In [16] an implementation was given for determining minimal sets of outputs that make a system structurally locally identifiable.

## 3 Theory

The definition of **ν**-identifiability and the justification of our results are made rigorous in Section 5. We give a brief summary of this theory here.

The notion of identifiability involves many subtleties. Our precise definition is a type of generic local single-experiment identifiability. This is done in terms of uniqueness (up to a finite set) of parameter and state values given known values of the output derivatives **y**^(≤**ν**)^ at a certain point in time. We show that if all derivatives of all outputs are available, then our definition is equivalent to the notion of local identifiability given in [10, Def. 2.5].

Our definition of **ν**-identifiability is intended to capture the property that a modeler really wants to ascertain, but it is often difficult to verify directly. Therefore we show that this property is equivalent to a property involving ranks of matrices, which is straightforward to check using existing software. This is in fact the property alluded to in Algorithm 1.

Although this rank property is straightforward to check, it involves computing ranks of symbolic matrices, which can cost the modeler time. Therefore, to alleviate this problem, we present probabilistic algorithms that substitute random integers for the variables and compute modulo a random prime. These are modifications of the method described in [8]. We present detailed probabilistic statements that a reader can use to create an implementation without investigating the proofs. We present detailed proofs that the reader can check without excessive pencil and paper computations.

Moreover, our work addresses an oversight in [8] surrounding the probability that a rational expression vanishes when evaluated at a random tuple of integers modulo a random prime. The author assumes that the denominator does not vanish, but then gives a bound for the probability that the numerator does not vanish regardless of this condition. We give a bound on the probability that the numerator does not vanish given that the denominator does not vanish.

## 4 Conclusions

Practical identifiability analysis lies at the heart of model-based data analysis. State-of-the-art numerical methods, such as profile likelihood and MCMC, suffer from the limitation that they require an exhaustive coverage of the parameter space, which is not possible for large and realistic models. This coverage is not needed for structural methods, but these methods traditionally assume that one can determine all derivatives of the measurements *y*. Such derivatives are not available in practice, but the consequences of this unfulfilled assumption have not been examined, because corresponding methods and algorithms are missing. Herein, we present a new definition – ν-identifability – which restricts the analysis to those derivatives of *y* that are available. We also present a new algorithm, which can determine such identifiability in practice. Applications to previously published models demonstrate that we can determine not only identifiability of parameters, but observability of any model property (Examples 2.4 and 2.5). These results allow us to strengthen previous conclusions. For instance, the previous rejection of a specific model for insulin signaling (Figure 2), was based on the model-guided experiments measuring the amount of internalized and dephosphorylated insulin receptors (Example 2.4), and we now show that this model property indeed can be identifiable, using a methodology that does not require coverage of the entire parameter space. Our analysis also shows that the number of parameters identifiable according to traditional structural methods are widely overestimated. If one e.g. assumes that only up to third order derivatives, instead of all derivatives, are available, the number of identifiable parameters drops from 17 to 1 for the Drosophila model (Figure 5), and from 21 to 4 for an NF-κB model (Figure 7). In both these models, the previously obtained identifiability is present only if at least 20 derivatives of all measurement signals are available. Our results bring us one step closer to a structural approach for practical identifiability analysis.

## 5 Technical Details

### 5.1 Notation and Definitions

The notion of identifiability involves many subtleties. This subsection provides the setup necessary for rigorous treatment.

For positive integers *n, ℓ, r*, and *m*, tuples **f** ∈ ℂ(**x, θ, u**)^*n*^ and **g** ∈ ℂ(**x, θ, u**)^*m*^, we have a model class

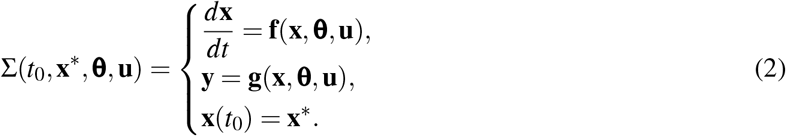

Fix positive integers *n, r*, and *m*, fix non-negative integer *ℓ*, and fix *n*-tuple **f** and *m*-tuple **g** of elements in ℂ(**x, θ, u**) for the rest of this section. The symbol ℕ denotes the set of non-negative integers.

#### Notation 5.1.

(Differential Algebraic Setting)

a. A *differential ring* (*S*, ^′^) is a ring together with a *derivation*, that is, a function ^′^ : *S* → *S* satisfying ∀*a, b* ∈ *S* (*a* + *b*)^′^ = *a*^′^ + *b*^′^ and (*ab*)^′^ = *a*^′^*b* + *ab*^′^. All rings are assumed to be commutative and with multiplicative identity. For *a* ∈ *S* and *k* ∈ ℕ, *a*^(*k*)^ denotes the image of *a* after *k* applications of ^′^. The set {*a*^(*k*)^ | *k* ≥ 0} will be called the set of *derivatives* of *a*.
b. Let (*S*, ^′^) be a differential ring and let *Z* = {*Z*_1_, …, *Z*_*k*_} be a set of indeterminates. Then *S*{*Z*} = *S*{*Z*_1_, …, *Z*_*k*_} denotes the polynomial ring in infinitely many indeterminates 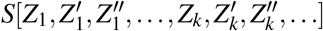 with derivation ^′^ extended by 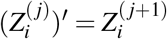.
c. Let (*S*_1_, ^′^) be a differential subring of (*S*_2_, ^′^) and let *Z* = {*Z*_1_, …, *Z*_*k*_}⊆ *S*_2_. Then *S*_1_ {*Z*} = *S*_1_ {*Z*_1_, …, *Z*_*k*_} denotes the smallest subring of *S*_2_ containing *S*_1_ and the derivatives of the elements of *Z*.
d. Let (*S*_1_, ^′^) be a differential ring that is a subring of a differential integral domain (*S*_2_, ^′^) and let *Z* = {*Z*_1_, …, *Z*_*k*_} ⊆ *S*_2_. Then *S*_1_*(Z)* = *S*_1_*(Z*_1_, …, *Z*_*k*_) denotes the fraction field of *S*_1_{*Z*}. This is a differential ring under the extension of ^*t*^ via the quotient rule (*a/b*)^′^= (*ba*^′^ −*ab*^′^)*/b*^2^.
e. For any differential ring containing any elements of ℂ(**θ**) we assume ^*t*^ maps each such element to 0.
f. Let (*S*, ^*t*^) be a differential ring. A *differential ideal I* is an ideal satisfying ∀*a* ∈ *I a*^′^ ∈ *I*. For subset *A* ⊆ *S*, we denote by [*A*] the smallest differential ideal containing *A*.
g. For an ideal *I* and element *a* in a ring *S*, we denote *I* : *a*^∞^ = {*s*∈ *S* |∃*k* : *a*^*k*^*s*∈ *I*}. This set is also an ideal in *S*.
h. Write **f** and **g** as fractions of elements of ℂ[**x, θ, u**], let *Q* ∈ ℂ[**x, θ, u**] be the LCM of the denominators, and let **F** and **G** be such that **f** = **F***/Q* and **g** = **G***/Q*. Note that *Q* is unique up to multiplication by an element of ℂ and choice of *Q* will not affect our definitions and results. We define the differential ideal of Σ as 𝒥 = [*Q***x**^′^− **F**, *Q***y** − **G**] : *Q*^∞^ ⊆ ℂ[**θ**]{**x, y, u**}. This is a prime (differential) ideal (cf [10, Lemma 3.2]) and thus *R* := ℂ[**θ**]{**u, x, y**}*/*𝒥 is a differential ring that is an integral domain.
i. For **ν** ∈ ℕ^*m*^, we denote by **y**^(≤**ν**)^ the subset 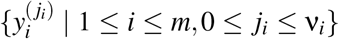 of *R*. We define ν_*max*_ := max(ν_1_, …, ν_*m*_).

#### Notation 5.2.

(Analytic Setting)

a. Let *t*_0∈_ ℂ. Then ℂ^∞^(*t*_0_) denotes the set of all functions that are complex analytic in some neighborhood of *t* = *t*_0_.
b. Let *t*_0_∈ ℂ. Then 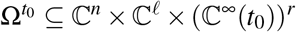 denotes the complement to the set where at least one of the denominators of **f** or **g** vanishes at *t* = *t*_0_.
c. For *t*_0_ ∈ ℂ and *h* ∈ ℂ(**x, θ**), let 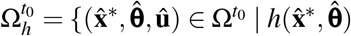 is defined}.
d. For *t*_0_ ℂ and 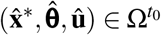,let 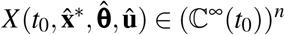 and 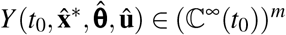 denote the unique solution of the initial value problem 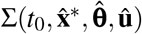 (cf. [24, Theorem 2.2.2]). For appropriate integers *j* and *k*, let 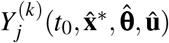 denote the *k*-th derivative of the *j*-th component of 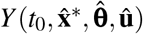.
e. For *s* ∈ℕ, a subset *U*⊆ ℂ^*s*^ is called *Zariski open* if there exists *P* in ℂ[*Z*_1_, …, *Z*_*s*_] such that *U* is the complement to the zero set of *P*.
f. For *s*∈ ℕ and *t*_0_ ∈ℂ, a subset *U*⊆ (ℂ^∞^(*t*_0_))^*s*^ is called *Zariski open* if there exists *P* in ℂ {*Z*_1_, …, *Z*_*s*_} such that

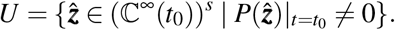
g. For *s* ∈ ℕ, *t*_0_∈ ℂ, and *W* = ℂ^*s*^ or (ℂ^∞^(*t*_0_))^*s*^, the set of all nonempty Zariski open subsets of *W* will be denoted by τ(*W*).
h. A subset *V* ⊆ ℂ is called *codiscrete* if for any distinct *c*_1_, *c*_2_ ∈ ℂ\*V* there exist disjoint open sets *W*_1_∋ *c*_1_ and *W*_2_ ∋*c*_2_ such that *W*_1_ ∩ (ℂ\*V*) = {*c*_1_} and *W*_2_ ∩ (ℂ\*V*) = {*c*_2_}.

#### Notation 5.3.

Let *h* ∈ ℂ(**x, θ**) ⟨ **u** ⟩, let *t*_1_ ∈ ℂ, and let 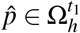.

- Substituting 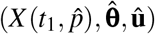 for (**x, θ, u**) in *h* gives a function of one variable. The image of *h* under this substitution will be denoted by 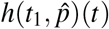.
- The connected component containing *t*_1_ of the intersection of the domains of 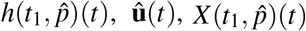 and 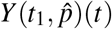 will be denoted by 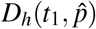.

#### Definition 5.4.

Let **ν** ∈ ℕ^*m*^. Let *h* ∈ ℂ(**x, θ**). The expression *h* is said to be **ν**-identifiable if

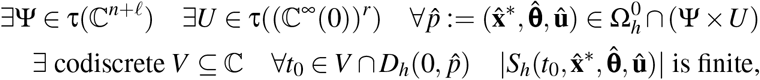

where

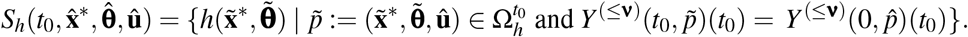

One can interpret this definition as follows: If at some time *t*_0_, not belonging to a certain discrete “singular” set, we know the exact values of *Y* ^(≤**ν**)^(*t*_0_), then if we consider all possible values of the state variables and parameters at *t*_0_ that produce these values of 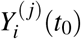,and then we consider the set of values of *h* obtained by evaluating *h* at these values, this set is finite.

#### 5.1.1 Examples to illustrate the definition

##### Example 5.5.

Consider the system

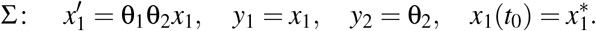

We show that θ_1_ is (1, 0)-identifiable. Note that *n* = 1 and *m* = 2. Since two parameters occur we can take *ℓ* to be any integer at least 2. Although no inputs occur the definitions require that we take *r* to be at least 1. Set ℓ= 2 and *r* = 1. Let 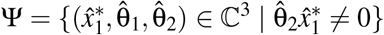 and let *U* = ℂ^∞^(0). Note that since there are no denominators we have 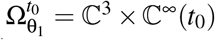 for all *t*_0_. Fix 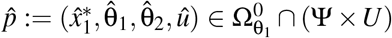. Let *V* = ℂ and fix 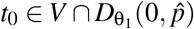.It is trivial to obtain a formula for the solution of this system, and we have

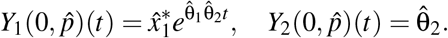

Let 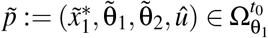,and note that

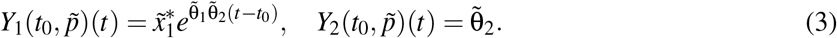

For **ν** = (1, 0), the condition 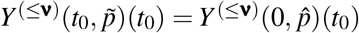 gives

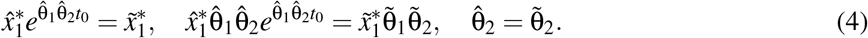

By the definition of Ψ, it follows that no side of the first or third of these equations is 0. Dividing the second equation by the product of the first and third proves that 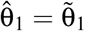.Since 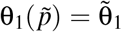,we see that 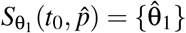 and thus has cardinality 1. We conclude that θ_1_ is (1, 0)-identifiable.

We now show that θ_1_ is not (0, 0)-identifiable. Fix Ψ ∈ τ(ℂ^3^) and *U* ∈ τ(ℂ^∞^(0)). Let 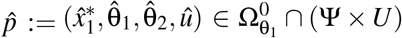. Fix codiscrete *V*, and let 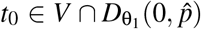,Using the first and third equations of (4), we see that for all *k* ∈ ℂ the tuple 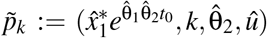 satisfies the condition 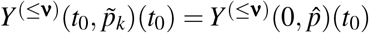 for **ν** = (0, 0). Therefore

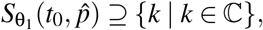

and the right-hand side is infinite. We conclude that θ_1_ is not (0, 0)-identifiable.

The next example includes an input and demonstrates why sometimes a discrete set must be excluded from *V*.

##### Example 5.6.

Consider the system

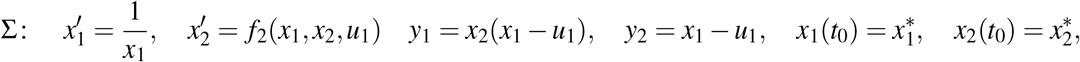

where *f*_2_ is some rational function.

We have *n* = 2 and *m* = 2. Take *ℓ* = 0 and *r* = 1. We show that *x*_2_ is (0, 0)-identifiable. We have 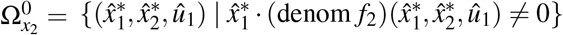. Let Ψ = ℂ^2^ and let *U* = {*z* ∈ ℂ^∞^(0) | *z*(0)*z* ^′^ (0) − 1 */*= 0}. Let 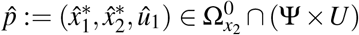. Note that because of the way *U* was defined, 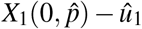 is not the zero function. Let *V* be the complement of the vanishing of 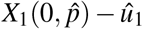,which is necessarily codiscrete since 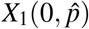 and û _1_ are analytic. Let 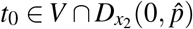. if 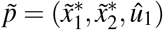 is such that 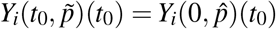 for *i* = 1, 2 then it follows that

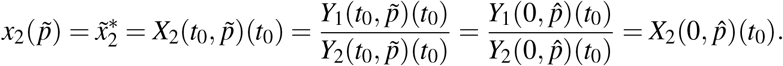

Since 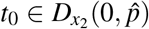 we know that 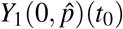 and 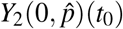 are defined and since *t*_0_ ∈ *V* we know that 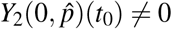.Thus 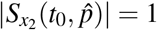 and we conclude that *x*_2_ is (0, 0)-identifiable.

#### 5.1.2 ν-identifiable implies single-experiment locally identifiable and the converse is true for sufficiently largeν

Many notions of identifiability are used in the literature. Some of these are stated and compared in [25]. In general, Definition 5.4 is not equivalent to any published definition as far as we are aware.

Definition 5.4 can be viewed as a type of single-experiment generic local identifiability. It is generic because there is a (possibly empty) set of numerical values of parameters and input functions for which *h* cannot be recovered but if the true values of the parameters and inputs lie outside this set *h* can be recovered. It is local because we allow *h* to lie in a finite set. One could change “is finite” to “equals one” to create a definition of a globally **ν**-identifiable function, however we do not address this in the current work. It is “single-experiment” because it refers to only one instance of the model. This is in contrast to a multiexperiment approach where one observes multiple instances of the model, usually with the same equation parameters but different initial conditions.

Since the equations of Σ are rational in **x, θ**, and **u**, there exist relations among sufficiently high derivatives of **y** and hence it is not meaningful to consider ν_*i*_ beyond a certain order. This is made precise by the following proposition.

##### Proposition 5.7.

*Let h* ∈ ℂ(**x, θ**) *and* **ν** ∈ ℕ^*m*^. *For each i* = 1, …, *m let µ*_*i*_ *be the greatest non-negative integer such that the set* 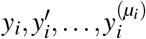 *is not algebraic over* ⟨ℂ**u**⟩. *Then h is* **ν***-identifiable if and only if h is* (min(ν_1_, *µ*_1_), …, min(ν_*m*_, *µ*_*m*_))*-identifiable. Moreover*, ∀*i µ*_*i*_ ≤ *n* + ℓ− 1.

*Proof*. Since the transcendence degree of ⟨ℂ**u**⟩ (**x, θ**) over ⟨ℂ**u**⟩is *n* + ℓ, it must be that 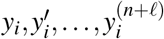 is algebraic over ⟨ℂ**u**⟩. Hence such a *µ*_*i*_ exists and moreover *µ*_*i*_ ≤*n* +− ℓ1.

For the ⇐ direction, note that if **a, b**∈ ℕ^*m*^ are such that *a*_*i≥*_ *b*_*i*_ for all *i*, it follows from the definition that *h* is **b**-identifiable implies *h* is **a**-identifiable.

We now address the ⇒ direction. Suppose *h* is **ν**-identifiable. By Proposition 5.12, *h* is algebraic over ⟨ℂ**u**⟩ (**y**^(≤**ν**)^). For any *i*, by writing an algebraic dependence of 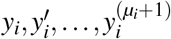 over ⟨ℂ**u**⟩ and differentiating (noting that the field has characteristic 0), we see that for all *s* > *µ*_*i*_ the element 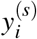 is algebraic over 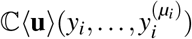.It follows that *h* is algebraic over 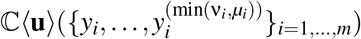.By Proposition 5.12 *h* is (min(ν_1_, *µ*_1_), …, min(ν_*m*_, *µ*_*m*_))-identifiable.

##### Definition 5.8.

Let *h*∈ ℂ(**x, θ**). The expression *h* is said to be single-experiment locally identifiable (SELI) if

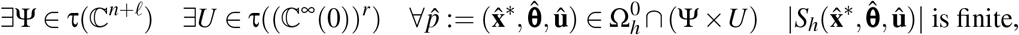

Where

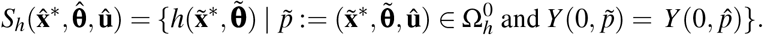

For *h* ∈ **x** ∪ **θ**, Definition 5.8 is equivalent to the definition of local identifiability given in Definition 2.5 of [10] (generalization to multiple inputs is asserted in Remark 2.2). In [10, Prop. 3.4 (a) ⇔ (c)], it was shown that for *h* ∈ **x** ∪ **θ**, *h* is SELI if and only if *h* is algebraic over ⟨ℂ**y, u**⟩. We extend this result to arbitrary *h* ∈ ℂ(**x, θ**).

##### Proposition 5.9.

*Let h* ∈ ℂ(**x, θ**). *Then h is SELI if and only if h is algebraic over* ℂ⟨**y, u**⟨.

*Proof*. Let 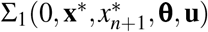 be the system obtained by adding to Σ(0, **x**^∗^, **θ, u**) the equations 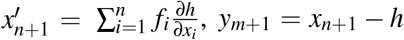, and 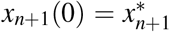.Note that 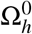 in Σ is the projection of 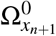 onto all coordinates but *x*_*n*+1_. We divide the proof into the following three steps: *h* is 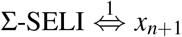 is 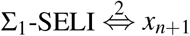 is algebraic over 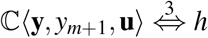 is algebraic over ⟨ ℂ**y, u**⟩.

We show *h* is Σ-SELI ⇔ *x*_*n*+1_ is Σ_1_-SELI. Suppose *h* is Σ-SELI. Let Ψ and *U* be as required by the definition. Define 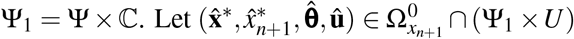. We verify that 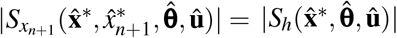 Since 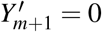,we have

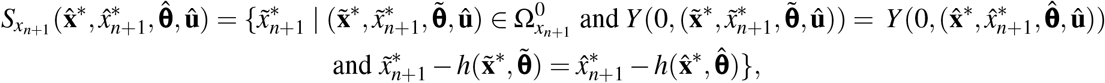

where *Y* = (*Y*_1_, …, *Y*_*m*_). It follows that

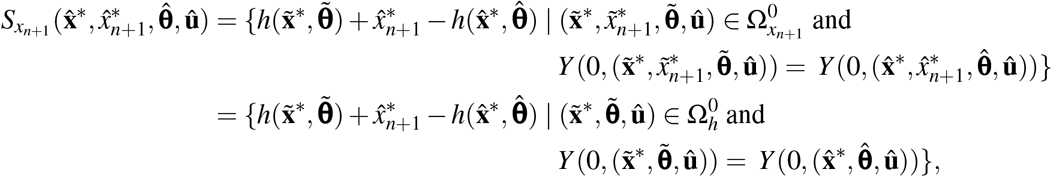

where 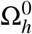 is considered a subset of ℂ^*n*+ ℓ^ *×* ℂ^∞^(0). Thus

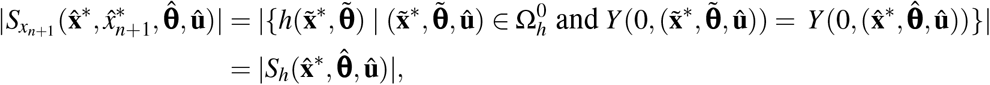

and we know that 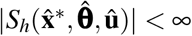 because *h* is Σ-SELI. This completes the first direction. Now suppose *x*_*n*+1_ is Σ_1_-SELI. Let Ψ_1_ and *U* be as required by the definition. Define Ψ to be the projection of Ψ_1_ onto all coordinates other than *x*_*n*+1_. Let 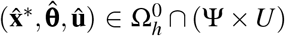.Let 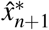 be such that 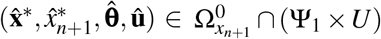.Such a 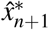 exists because of the way Ψ was defined. Now

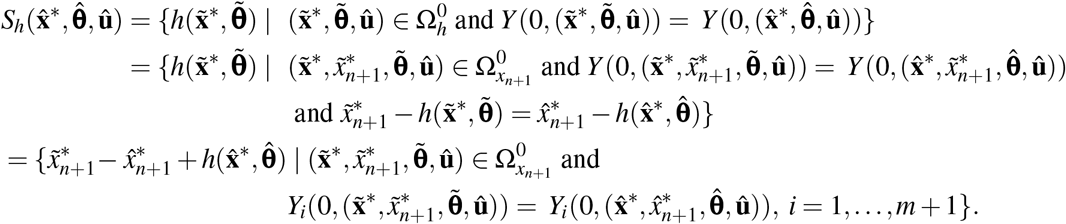

Hence

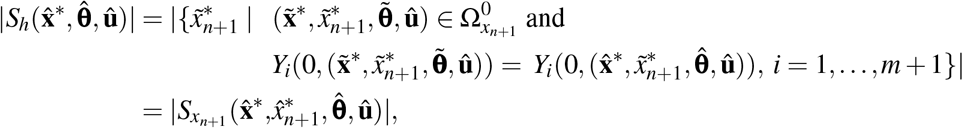

and we know that 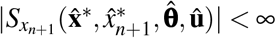 because *x*_*n*+1_ is Σ_1_-SELI. ⟨

From [10, Prop. 3.4 (a) ⇔ (c)], we have that *x*_*n*+1_ is Σ_1_-SELI ⇔ *x*_*n*+1_ is algebraic over ℂ(**y**, *y*_*m*+1_, **u**). We now show that *h* is algebraic over ⟨ℂ**y, u**⟩ ⇔*x*_*n*+1_ is algebraic over ℂ⟨**y**, *y*_*m*+1_, **u** ⟩. Suppose *h* is algebraic over ℂ⟨**y, u**⟩. Then *x*_*n*+1_ = *h* + *y*_*m*+1_ is algebraic over ℂ⟨**y**, *y*_*m*+1_, **u**⟩. We now address the other direction. Let ℱ = ℂ(**y, u**) and let ℱ_1_ = ℂ⟨**y**, *y*_*m*+1_, **u**⟩. Suppose *x*_*n*+1_ is algebraic over ℱ_1_. Then *h* = *x*_*n*+1_ − *y*_*m*+1_ is algebraic over ℱ_1_. Let 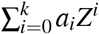 be the minimal polynomial of *h* over ℱ_1_. Suppose *y*_*m*+1_ appears in some *a*_*i*_. Let α be the differential field automorphism on ℂ(**x**, *x*_*n*+1_, **θ**) ⟨**u**⟩ such that ℂ(**x, θ**) ⟨**u**⟩ is fixed pointwise, α(*y*_*m*+1_) = *y*_*m*+1_ + 1, and α(*x*_*n*+1_) = *x*_*n*+1_ + 1. Now 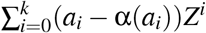 is a non-zero polynomial of lower degree with coefficients in ℱ_1_ that has *h* as a root. Thus we have a contradiction. Therefore all the *a*_*i*_ lie in ℱ and *h* is algebraic over ℱ.

This leads to the main result of this section.

##### Corollary 5.10.

*Let h* ∈ ℂ(**x, θ**) *and* **ν** ∈ ℕ^*m*^.

1. *h is* **ν***-identifiable* ⇒ *h is SELI*.
2. *If* ∀*i* ν_*i*_ ≥ *n* + *ℓ* − 1, *then h is SELI* ⇒ *h is* **ν***-identifiable*.

*Proof*. Suppose *h* is **ν**-identifiable. Then by Proposition 5.12 we know *h* is algebraic over ℂ⟨**u**⟩ (**y**^(≤**ν**)^). It follows that *h* is algebraic over ℂ⟨ **y, u**⟩ and then by Proposition 5.9 we have that *h* is SELI.

Suppose that ∀*i* ν_*i*_ ≥ *n* + *ℓ* − 1. Then by the arguments presented in Proposition 5.7 we have ℂ⟨**u**⟩ (**y**^(≤**ν**)^) = ℂ⟨ **y, u**⟩. Suppose *h* is SELI. By Proposition 5.9 we have that *h* is algebraic over ℂ⟨ **y, u**⟩, which equals ℂ⟨**u**⟩ (**y**^(≤**ν**)^). By Proposition 5.12 we have that *h* is **ν**-identifiable.

### 5.2 Proof of Algorithms

The definition of **ν**-identifiability is stated in terms of analytic functions. The Proposition 5.12 gives a correspondence between the analytic property and an algebraic property. Its proof will use the following lemma.

#### Lemma 5.11.

*Let t*_1_,*t*_2_ ∈ ℂ, *and let*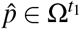. *The map*

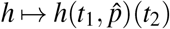

*gives a* ℂ*-algebra homomorphism S* → ℂ, *where S is the ring* 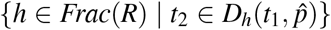.*Moreover, under this* 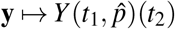.

*Proof*. Denote the stated map by φ. Since 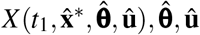 satisfy all the relations among **x, θ, u** and no denominator of *S* is sent to 0, we see that φ is a homomorphism on *S*. Now **y** = **g**(**x, θ, u**), so 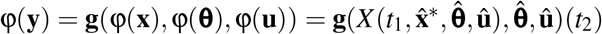.The existence and uniqueness theorem guarantees the existence and uniqueness of 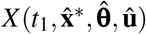,and 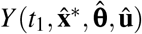 is given by 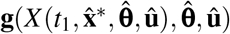. □

#### Proposition 5.12.

*Let* **ν** ∈ ℕ^*m*^. *Expression h* ∈ ℂ(**x, θ**) *is* **ν***-identifiable if and only if h is algebraic over the subfield* ℂ⟨**u**⟩ (**y**^(≤**ν**)^) *of Frac*(*R*).

*Proof*. Part 1: ⇐ Assume that *h* is algebraic over ℂ⟨**u**⟩ (**y**^(≤**ν**)^). Consider the minimal polynomial of *h* over ℂ⟨**u**⟩ (**y**^(≤**ν**)^). Clearing denominators, we obtain the polynomial 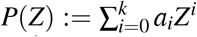,where each *a*_*i*_ ∈ ℂ{**u**}[**y**^(≤**ν**)^] and *P*(*h*) = 0. By [26, Corollary 6.6], there exist Ψ ∈ τ(ℂ^*n*+*ℓ*^) and *U* ∈ τ((ℂ^∞^ (0))^*r*^) such that for all 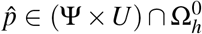 it holds that 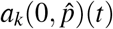 (see Notation 5.3) is not the zero function. Fix such Ψ and *U* and choose a 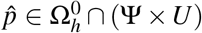.Let *V* be the complement of the vanishing of *a* 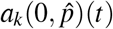.Since 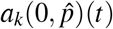 is analytic about 0, we know that *V* is codiscrete.

Fix 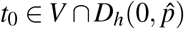,Let 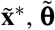 be such that 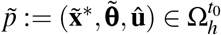 and 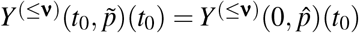. Since each *a*_*i*_ belongs to ℂ{**u**}[**y**^(≤**ν**)^], it follows from Lemma 5.11 that 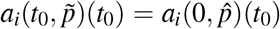 for all *i*. Applying Lemma 5.11 to the equation *P*(*h*) = 0, we find that 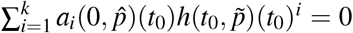.Thus 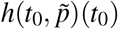 is a root of the non-zero polynomial 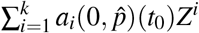. Noting that 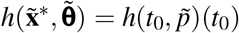 we conclude that 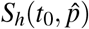 is finite.

Part ⇒ 2: Assume that *h* is not algebraic over ℂ ⟨**u**⟩ (**y**^(≤**ν**)^) and that *h* is **ν**-identifiable. We will give a proof by contradiction. The proof can be divided into four steps:

**Step 0** Label the rings that will be used in the proof.

**Step 1** Choose 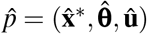.This will be done in terms of the non-vanishing of minimal polynomials over fraction fields of intermediate rings. Fix *V* and *t*_0_ and note that Lemma 5.11 with 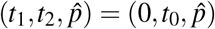 gives a ring homomorphism φ: *R*[*h*] → ℂ.

**Step 2** Show that the set δ(*h*) δ: *R*[*h*] → ℂ is a ℂ-algebra homomorphism such that δ(**u**^(ℕ)^) = φ(**u**^(ℕ)^) and δ(**y**^(≤**ν**)^) = φ(**y**^(≤**ν**)^) is infinite. This will involve careful extension of ring homomorphisms.

**Step 3** Verify that 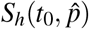 is infinite by noting that each δ corresponds to a tuple 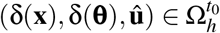.

**Step 0** Label *x*_1_, …, *x*_*n*_, θ_1_, …, θ_*ℓ*_ as *b*_1_, …, *b*_*n*+*ℓ*_. Without loss of generality, let *k* be such that *j≤ k* if and only if *b* _*j*_ is algebraic over ℂ ⟨**u**⟩ (**y**^(≤**ν**)^). Let *R*_2_ equal ℂ {**u}** [**y**^(≤**ν**)^, *b*_1_, …, *b*_*k*_] and let *F*_2_ be its field of fractions. Let *µ* be such that *b*_*k*+1_, …, *b*_*µ*_, *h* is a transcendence basis for Frac(*R*) = ℂ ⟨**u**⟩ (**x, θ**) over *F*_2_, relabeling if necessary. Let *R*_3_ = *R*_2_[*b*_*k*+1_, …, *b*_*µ*_] and let *F*_3_ be its field of fractions. Now *b*_*µ*+1_, …, *b*_*n*+*ℓ*_ are each algebraic over *F*_4_ := *F*_2_(*b*_*k*+1_, …, *b*_*µ*_, *h*). For *j* = *µ* + 1, …, *n* + *ℓ* let *P*_*j*_(*Z*) be the minimal polynomial of *b* _*j*_ over *F*_4_ multiplied by the LCM of the denominators, where *F*_4_ is viewed as the field of fractions of *R*_4_ := *R*_2_[*b*_*k*+1_, …, *b*_*µ*_, *h*]. For each *j* = *µ* + 1, …, *n* + *ℓ*, let *L*_*j*_(*b*_*k*+1_, …, *b*_*µ*_, *h*) ∈ *R*_4_ be the leading coefficient of *P*_*j*_. Write *h* = *n*_*h*_*/d*_*h*_, where *n*_*h*_, *d*_*h*_ ∈ *R*. Let 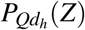 be the minimal polynomial of *Q*· *d*_*h*_ over *F*_4_ multiplied by the LCM of the denominators. (Note that if *Qd*_*h*_ ∈ *F*_4_ then 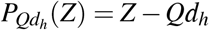.Let 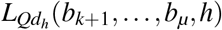 and 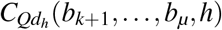 be its leading and constant coefficients, respectively.

**Table 1:**
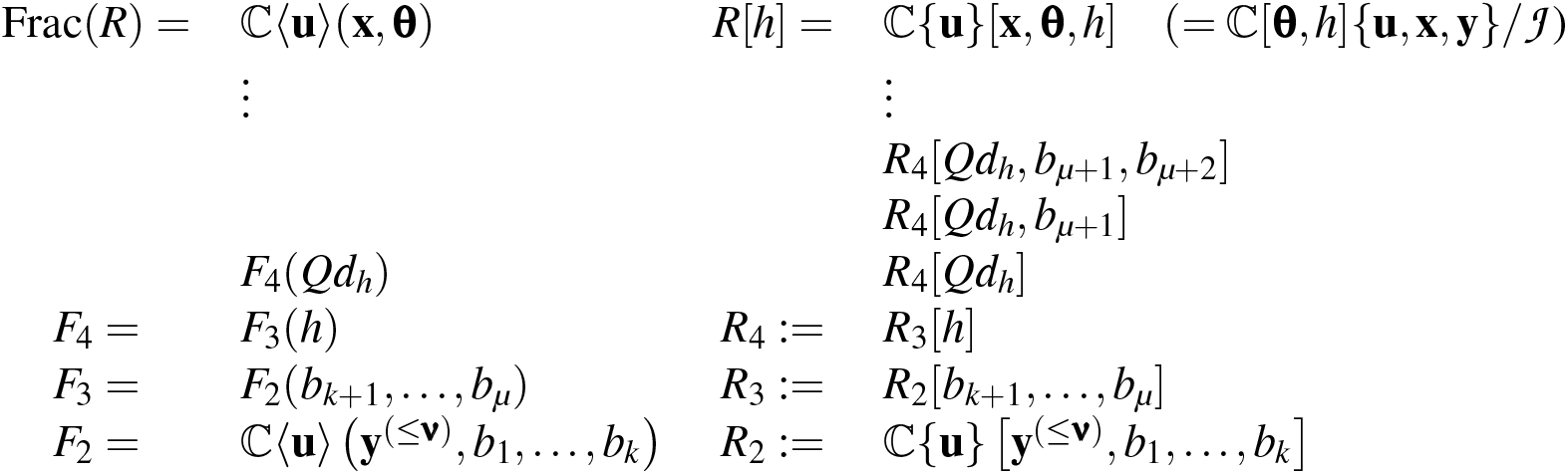
Intermediate rings and their fraction fields.

**Step 1** Let Ψ and *U* be as guaranteed by the definition of **ν**-identifiability. Consider the subset of 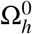 consisting of all 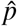 such that for all *j* = *µ* + 1, …, *n* + *ℓ* 1 the expressions 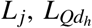,and 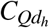 evaluated at 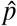 and then *t* = 0 are not zero. This subset is non-empty because it is the intersection of finitely many non-empty Zariski open sets. Hence its intersection with Ψ ×*U* is non-empty. Fix such a 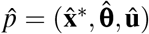.Let *V* be as required by the definition of **ν**-identifiability, let 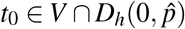,(recall Notation 5.3), and let φ be the ℂ-algebra homomorphism from *R*[*h*] to ℂ given by applying the ring homomorphism described in Lemma 5.11 with 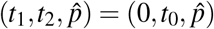.

**Step 2** We now show that there are infinitely many *c* ∈ℂ such that there exists a ℂ-algebra homomorphism δ: *R*[*h*] →ℂ such that δ(**u**^(ℕ)^) = φ(**u**^(ℕ)^), δ(**y**^(≤**ν**)^) = φ(**y**^(≤**ν**)^), and δ(*h*) = *c*.

Define δ on *R*_2_ by δ := φ | _*R*2_. For each *j*, define δ(*L*_*j*_)(*b*_*k*+1_, …, *b*_*µ*_, *h*) ∈ ℂ[*b*_*k*+1_, …, *b*_*µ*_, *h*] to be the element obtained by evaluating the coefficients (in *R*_2_) of *L*_*j*_ via δ; define the analogous expressions with *L*_*j*_ replaced by 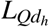 and 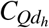. By the way 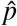 was chosen, we have that 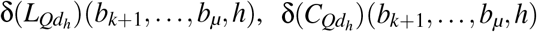, and all δ(*L*_*j*_)(*b*_*k*+1_, …, *b*_*µ*_, *h*) are non-zero. Using Lemma 5.13 we can extend δ to *R*_3_ := *R*_2_[*b*_*k*+1_, …, *b*_*µ*_] in a way that makes neither 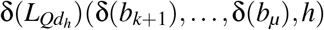 nor 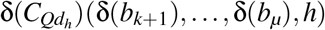 nor any δ(*L*_*j*_)(δ(*b*_*k*+1_), …, δ(*b*_*µ*_), *h*) ∈ℂ[*h*] equal to zero. Choose such an extension and call this, somewhat abusing notation, δ. Since *h* is not algebraic over *F*_3_, by Lemma 5.13 δ can be extended by mapping *h* to any element of ℂ. Now there are infinitely many *c* such that 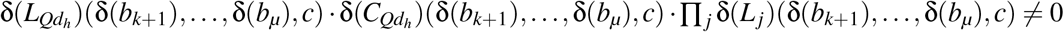. Fix such a *c* for the remainder of the proof and extend δ to *R*_4_ by letting δ(*h*) ≠*c*.

We now extend δ to *R*_4_[*Qd*_*h*_]. By the preceding discussion, the leading and constant coefficients of 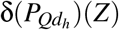 are non-zero, so by Lemma 5.13 we can extend δ to *R*_4_[*Qd*_*h*_] so that δ(*Qd*_*h*_) ≠0.

Next we extend δ to *R*_4_[*Qd*_*h*_, *b*_*µ*+1_]. Observe that the minimal polynomial *Q*_*µ*+1_(*Z*) of *b*_*µ*+1_ over *F*_4_(*Qd*_*h*_) is a factor of *P*_*µ*+1_(*Z*). Therefore δ(*Q*_*µ*+1_)(*Z*) is a non-zero factor of δ(*P*_*µ*+1_)(*Z*). By Lemma 5.13 we can extend δ to *R*_4_[*Qd*_*h*_, *b*_*µ*+1_].

The argument from the preceding paragraph can be repeated to show that δ can be extended to *R*_4_[*Qd*_*h*_, *b*_*µ*+1_, *b*_*µ*+2_]. Applying this several more times, we see that we can extend δ to *R*[*h*].

Let *T* = {δ |δ: *R*[*h*] →ℂ is a ℂ-algebra homomorphism such that δ(**u**^(ℕ)^) = φ(**u**^(ℕ)^) and δ(**y**^(≤**ν**)^) = φ(**y**^(≤**ν**)^)}. We have shown that {δ(*h*) | δ ∈*T*} is infinite.

**Step 3** We will now show that 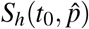 is infinite. We have

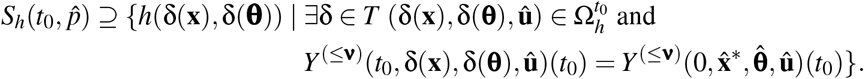

By Lemma 5.11 it holds that *Y* ^(≤**ν**)^(*t*_0_, δ(**x**), δ(**θ**), **û**)(*t*_0_) = δ(**y**^(≤**ν**)^) and 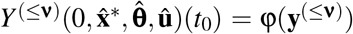.Hence

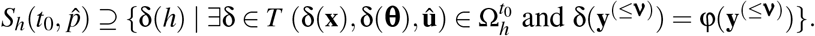

Since for each δ ∈ *T* we have δ(*Qd*_*h*_) ≠0, the first conjunct is satisfied. By the definition of *T*, the second conjunct is satisfied for all δ ∈ *T*. Hence

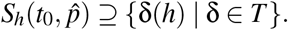

We showed in Step 2 that the right hand side is infinite, and thus 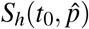 is infinite. Therefore *h* is not **ν**-identifiable, contradicting our assumption.

#### Lemma 5.13.

*Let W be a (possibly infinite) set of indeterminates and let S be a* ℂ*-subalgebra of the field of rational functions* ℂ(*W*). *Let* φ: *S*→ ℂ *be a* ℂ*-algebra homomorphism. Let f* ∈ℂ(*W*)\ {0}.

*Suppose f is not algebraic over Frac*(*S*). *Then* φ *can be extended to S*[*f*] *by mapping f to any element of* ℂ.

*Suppose f is algebraic over Frac*(*S*). *Let P*(*Z*) *be the minimal polynomial of f over Frac*(*S*) *multiplied by the LCM of the denominators. Write P*(*Z*) = *a*_*k*_*Z*^*k*^ + … + *a*_0_, *where a*_*k*_, …, *a*_0_ ∈ *S. If* φ(*a*_*k*_) ≠ 0, *then* φ *can be extended to S*[*f*]. *If furthermore* φ(*a*_0_) ≠ 0, *then in this extension* φ(*f*) ≠ 0.

*Proof*. Suppose *f* is not algebraic over Frac(*S*). Our result follows from [27, p. 99].

Suppose *f* is algebraic over Frac(*S*) and φ(*a*_*k*_) ≠ 0. If *f*∈ Frac(*S*) then the result is trivial. Assume *f* is not in Frac(*S*). By [27, Theorem 3.2 p. 347], φ can be extended to *S*[*f*] or *S*[*f* ^−1^]. If the former is true the proof is complete. Suppose φ can be extended to φ_1_ : *S*[*f* ^−1^] → ℂ. Writing 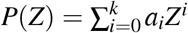,we have that 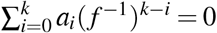. If φ_1_(*f* ^−1^) = 0, then it follows that 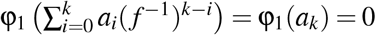, which contradicts our hypotheses. Thus φ_1_(*f* ^−1^) ≠ 0. Now we can extend φ_1_ to the ring 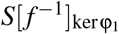,which is equal to the subring of ℂ(*W*) consisting of fractions with numerators in *S*[*f* ^−1^] and denominators in *S*[*f* ^−1^]\ kerφ_1_ (cf. [27, p. 346]). Since *f* ^−1^ ∉ kerφ_1_, the element *f* belongs to 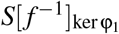. By restricting φ_1_ to *S*[*f*] we have an extension of φ to *S*[*f*].

Still assuming *f* is algebraic over Frac(*S*), suppose φ(*a*_0_) ≠ 0. The image of *f* must satisfy 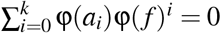. Thus it is impossible that φ(*f*) = 0.

It is not always obvious whether a given field element is algebraic over a given subfield. The following proposition gives an equivalence that, for our purposes, reduces this question to the problem of checking the rank of a matrix with easily computable entries. We will use the following notation:

#### Notation 5.14.

Let *W* = *W*_1_, …, *W*_*a*_ be elements of a ℂ-algebra and let *Z* = *Z*_1_, …, *Z*_*b*_ be an algebraically independent set over ℂ. We denote by 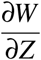 the matrix whose (*i, j*)-th entry is 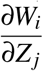,If *M* is such a matrix and *Z*_0_ ∈ *Z*, then *M\Z*_0_ denotes the result when the column corresponding to *Z*_0_ is removed. If this column does not appear in *M* then *M\Z*_0_ is equal to *M*.

We will use the following algebraic fact, which generalizes [28, Thm. 2.3].

#### Proposition 5.15.

*Let p*_1_, …, *p*_*s*_ *each be algebraic over* ℂ(*w*_1_, …, *w*_*a*_, *z*_1_, …, *z*_*b*_), *where s*≤ *b and all w*_*i*_ *and z*_*i*_ *are indeterminates. The elements p*_1_, …, *p*_*s*_ *are algebraically independent over* ℂ(*w*_1_, …, *w*_*a*_) *if and only if the matrix* ∂(*p*_1_, …, *p*_*s*_)*/*∂(*z*_1_, …, *z*_*b*_) *has rank equal to s*.

*Proof*. Let *M* := ∂(*p*_1_, …, *p*_*s*_)*/*∂(*z*_1_, …, *z*_*b*_) and let *M*_1_ := ∂(*w*_1_, …, *w*_*a*_, *p*_1_, …, *p*_*s*_)*/*∂(*w*_1_, …, *w*_*a*_, *z*_1_, …, *z*_*b*_).

Note that

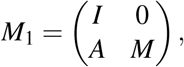

where *I* is the *a ×a* identity matrix and *A* is an *s× a* matrix.

Now *p*_1_, …, *p*_*s*_ is algebraically independent over ℂ(*w*_1_, …, *w*_*a*_) iff *w*_1_, …, *w*_*a*_, *p*_1_, …, *p*_*s*_ is algebraically independent over ℂ iff (by [28, Thm. 2.3]) rank *M*_1_ = *a* + *s* iff rank *M* = *s*.

#### Proposition 5.16.

*Let* **ν** ∈ ℕ^*m*^ *and let h* ∈ ℂ(**x, θ**). *Then h is algebraic over* ℂ⟨**u**⟩ (**y**^(≤**ν**)^) *if and only if the matrix* 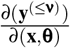 *has the same rank as* 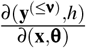.

*Proof*. Note that a subset of **y**^(≤**ν**)^ ∪{*h*} is algebraically independent over ℂ⟨**u**⟩ if and only if it is algebraically independent over 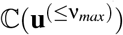.Let *T* be a maximal subset of **y**^(≤**ν**)^ that is algebraically independent over ℂ⟨**u**⟩. Then by Proposition 5.15 with 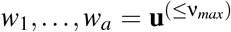 and *z*_1_, …, *z*_*b*_ = **x** ∪ **θ** we have 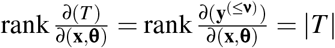.

Suppose *h* is algebraic overℂ⟨**u**⟩)(**y**^(≤**ν**)^). Then *T* ∪ {*h*} is algebraic over 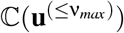. Then 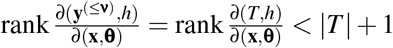 by Proposition 5.15. Therefore 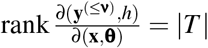.

Suppose *h* is not algebraic over ℂ⟨**u**⟩ (**y**^(≤**ν**)^). Then *T* ∪ {*h*} is not algebraic over 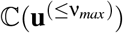. Then 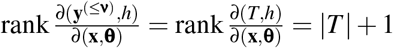 by Proposition 5.15. □

#### Corollary 5.17.

*Algorithm 1 always terminates. The output is “Yes” if and only if h is* **ν***-identifiable. Proof*. Termination is obvious. The other result follows from Proposition 5.12 and Proposition 5.16. □

#### Proposition 5.18.

*Let* **ν** ∈ ℕ^*m*^ *and let J be as in Algorithm 2. Let z* ∈ **x** ∪ **θ**. *Then z is algebraic over*ℂ⟨**u**⟩ (**y**^(≤**ν**)^) *if and only if* rank(*J/z*) = rank(*J*) − 1.

*Proof*. Without loss of generality we assume *z* = θ_*ℓ*_. By Proposition 5.16, θ_*ℓ*_ is algebraic over ℂ⟨**u**⟩ (**y**^(≤**ν**)^) if and only if rank 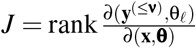.Now

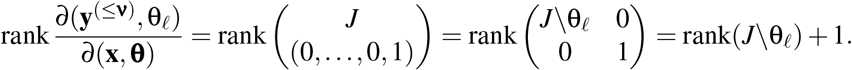

#### Corollary 5.19.

*Algorithm 2 always terminates. The output is “Yes” if and only if* θ_*i*_ *is* **ν***-identifiable. Proof*. Termination is obvious. The other result follows from Proposition 5.12 and Proposition 5.18. □

### 5.3 Probabilistic method for improved speed

#### 5.3.1 Presentation of algorithms and summary of results

Algorithms 1 and 2 involve computing the rank of a matrix of rational expressions. It is usually much faster to insert random numbers for the variables and compute the rank of the resulting matrix. The disadvantage of this is that the numerical matrix may have lower rank than the symbolic. In [29] a method for doing this in the case where the coefficients of Σ are integers with user-specified probability of success was given. We adapt that method to our algorithms.

For the rest of this section, assume the coefficients of **f** and **g** in (2) and *h* in Algorithm 1 are integers. Note that the entries of *J* belong to 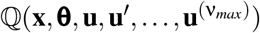. Our strategy involves randomly choosing *n* + *ℓ* + (ν_*max*_ + 1)*r* non-negative integers and a prime number, and then evaluating the determinant of our matrix at these integers modulo the prime. Algorithm 3 implements this strategy on Algorithm 1.

##### Algorithm 3

Determines whether parameter combination *h* is **ν**-identifiable

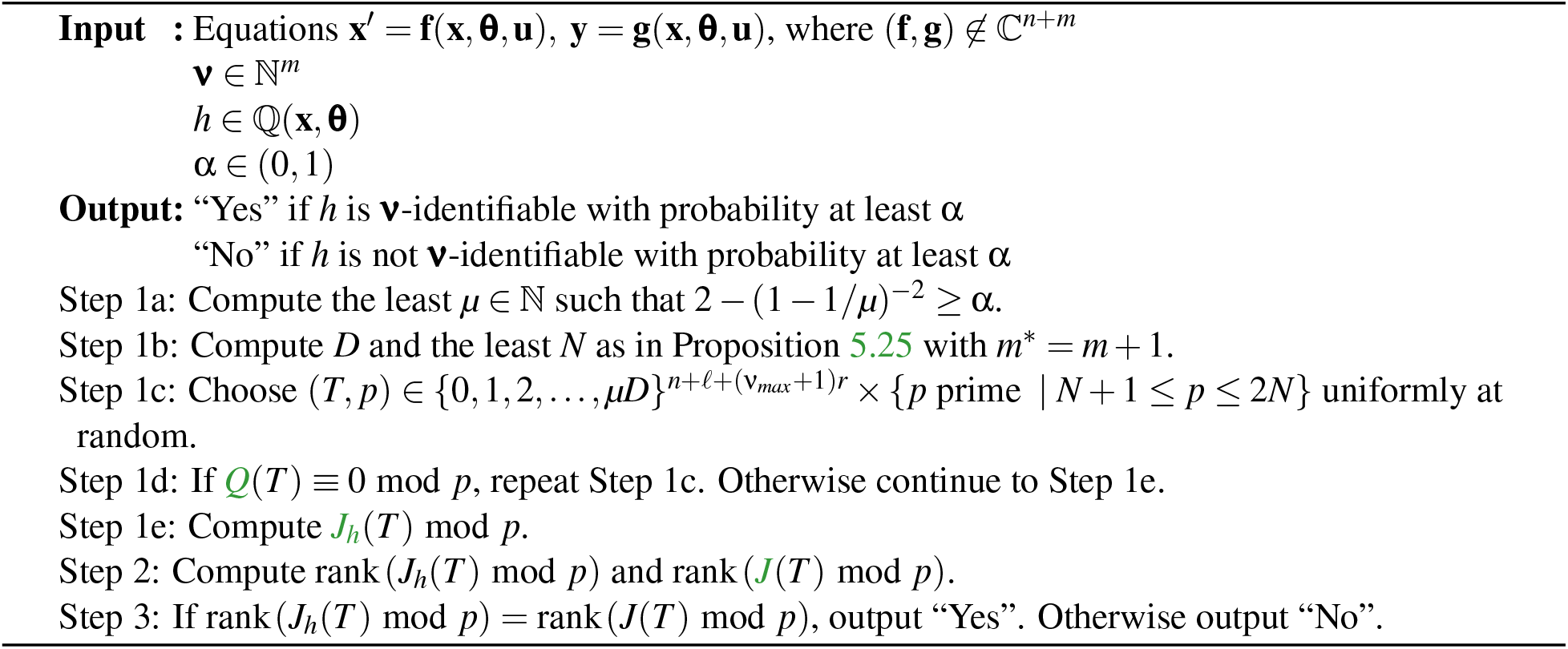

While Algorithm 2 determines whether an individual parameter is **ν**-identifiable, Algorithm 4 uses this concept to determine, with user-specified probability, all elements of **x** ∪ **θ** that are **ν**-identifiable.

##### Algorithm 4

Determines the **ν**-identifiable subset of **x** ∪ **θ**

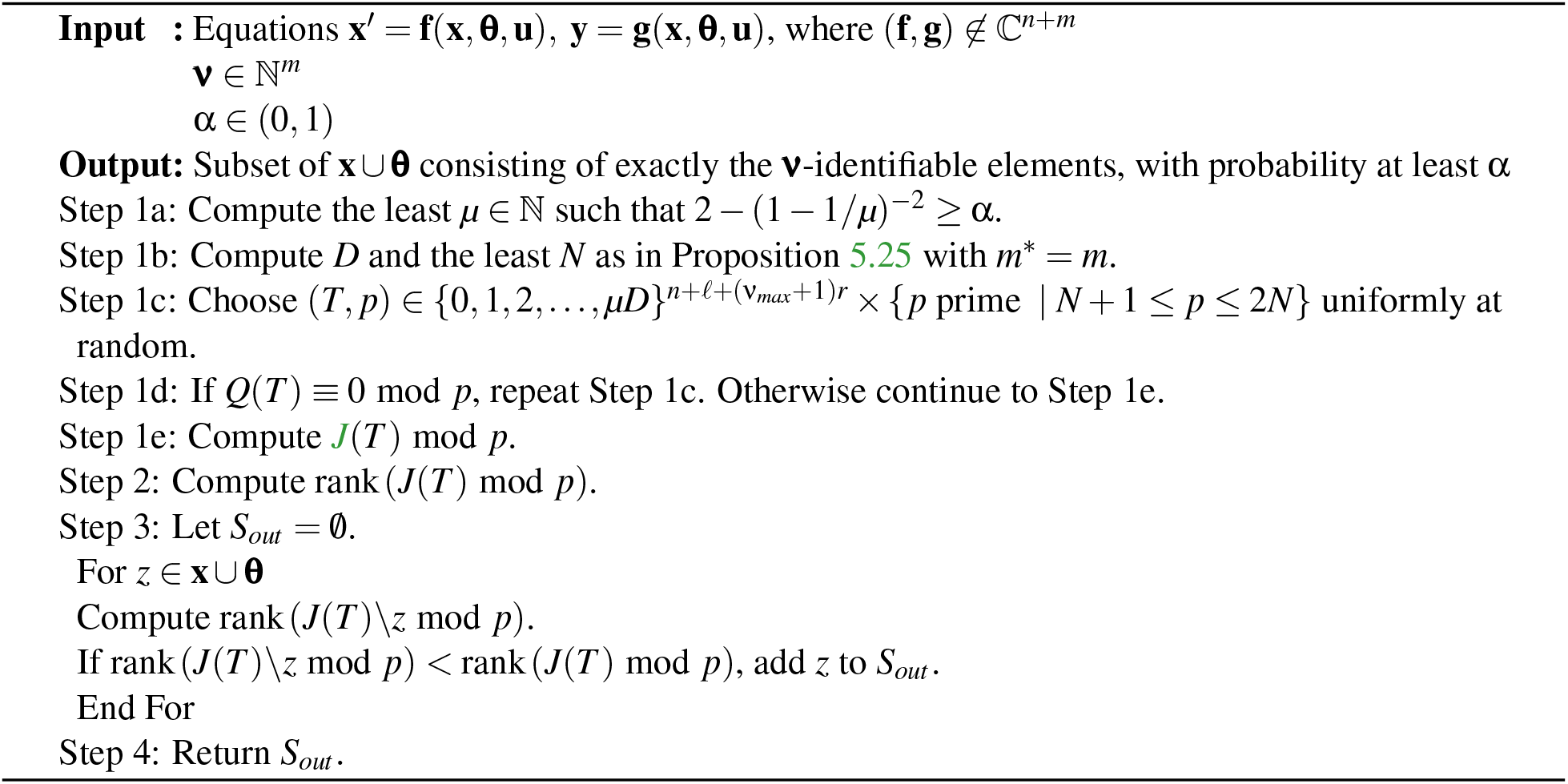

The main results on Algorithm 3 and Algorithm 4 are the following:

1. The expected time to reach Step 1e is negligible. Once Step 1e is reached the algorithm is guaranteed to terminate. (Proposition 5.29)
2. The probability that the output is correct is at least α. (Propositions 5.30 and 5.31)
3. For Algorithm 3, if in Step 2 rank(*J*(*T*) mod *p*) = *n* + *ℓ*, then *h* is **ν**-identifiable. For Algorithm 4, if in Step 2 rank(*J*(*T*) mod *p*) = *n* + *ℓ*, then every element of **x** ∪ **θ** is **ν**-identifiable. (Proposition 5.32)

##### Remark 5.20.

At the beginning of this section, we assumed *n, m*, and *r* are positive integers. As noted in Example 5.5, *ℓ* and *r* must be at least equal to the number of parameters and outputs, respectively, that appear in Σ, but can be chosen to be greater without changing identifiability results. In Algorithms 3 and 4 it is sufficient to use *ℓ* and *r* equal to the number of parameters and inputs, respectively, appearing in the equations. In particular one can use *r* = 0 if no inputs appear and the main results on the algorithms are correct.

##### Remark 5.21.

We have presented Algorithms 3 and 4 only for the case where **f** and **g** are not all elements of ℂ, since removing this restriction would require addressing special cases in several of the proofs. If (**f, g**) ∈ ℂ^*n*+*m*^, then *h* is **ν**-identifiable if and only if *h* ∈ ℂ. One could easily add a step at the beginning of either algorithm to accommodate this case.

##### Remark 5.22.

Our algorithms do not specify the methods used to compute the ranks of matrices. One can use the state-of-the-art method for this to achieve maximum speed.

#### 5.3.2 Proof of algorithms

In Algorithms 3 and 4, a random tuple of integers *T* and a random prime *p* are chosen, we check that the denominator of a determinant evaluated at *T* does not vanish modulo *p*, and then calculate the rank of a matrix after evaluating at *T* modulo *p*. We show that this gives the correct results with user-specified probability. The main results on Algorithms 3 and 4 are Propositions 5.29, 5.30, 5.31, and 5.32. The theory is based on bounds on integer roots of polynomials with integer coefficients, as well as the distribution of the prime numbers.

First, we use Proposition 5.23, Lemma 5.24, and Lemma 5.26 to prove Proposition 5.25, which gives conditions on the sets from which we choose *T* and *p* so that the numerator of the determinant evaluated at *T* does not vanish modulo *p* with user-specified probability. This proposition is essentially a more precisely stated version of [29, Proposition 6], and the proof we give is outlined in [29].

Next, Proposition 5.28 shows that the probability that the numerator vanishes given that the denominator does not vanish can be specified by the user. Note that this is the true probability associated with the algorithm, since we must first check that our choice of (*T, p*) does not make the denominator vanish before proceeding. This issue is not addressed in similar algorithms (cf. [8, p. 739], [21]) and is non-trivial, as shown by Example 5.27.

Finally, we prove statements about the algorithms that are directly relevant to helping the user interpret them. Although in principle, arbitrarily many instances of (*T, p*) may need to be chosen before finding one that does not make the denominator vanish, Proposition 5.29 shows that the expected time for a successful choice is negligible. It also asserts the algorithm’s termination after such a successful choice. Propositions 5.30 and 5.31 show that the algorithms produce the correct result with user-specified probability. Proposition 5.32 states that when the rank of the specialized matrix is full, the algorithms output the correct result with certainty.

##### Proposition 5.23

([30] Prop. 98 p. 192). *Let P*(*Z*_1_, …, *Z*_*k*_) *be a polynomial of total degree D over an integral domain A. Let S* ⊆ *A. If an element* (*z*_1_, …, *z*_*k*_) *is chosen from S*^*k*^ *uniformly at random, then*

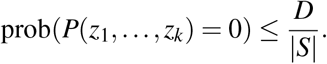

##### Lemma 5.24

([31] Lemma 18.9 p. 525). *Let S* ⊆ ℕ *be a nonempty finite set of prime numbers, let M* ∈ ℤ*\*{0}, *and let C* ≥ |*M*|. *If p is chosen from S uniformly at random, then*

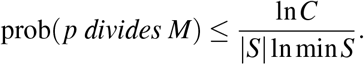

##### Proposition 5.25.

*Let J (resp. J*_*h*_*) be as in Algorithm 2 (resp. Algorithm 1) and suppose J (resp. J*_*h*_*) has at least one non-zero entry. Let*

1. *µ* ∈ ℕ^+^
2. *d*_1_ = *maximum degree of the numerators and denominators of* **f** *and* **g** *(resp*. **f, g** *and h)*
3. *d*_2_ = max{ln(|*a*| + 1) : *a is a coefficient in* **f** *or* **g** (*resp*. **f, g**, *or h*)}
4. *D* = (*n* + *ℓ*)(2ν_*max*_ + 3)(*n* + *m*^∗^)*d*_1_, *where m*^∗^ = *m (resp. m*^∗^ = *m* + 1)
5. *C be such that* ln*C* = (2 ln(*n* + *ℓ* + *r* + 1) +ln(*µD* + 1))*D* +(*n* + *ℓ*)(2ν_*max*_ + 3)((*n* + *m*^∗^)*d*_2_ +ln(2*nD*)), *where m*^∗^ = *m (resp. m*^∗^ = *m* + 1)
6. *N* ∈ ℕ *such that N* ≥ 2*µ* ln*C*
7. *S* = {*p prime* | *N* + 1 ≤ *p* ≤ 2*N*}

*Let J*_0_ *be a square submatrix of J (resp. J*_*h*_*) with rank equal to that of J (resp. J*_*h*_*). Note that the numerator of* det *J*_0_ *lies in* 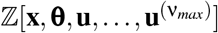.*If a tuple of values T of the variables is chosen uniformly at random from* 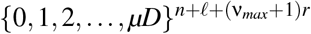 *and p is chosen uniformly at random from S, then*

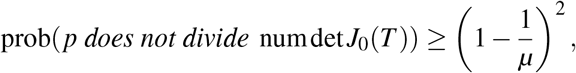

*where* numdet *J*_0_(*T*) *represents the specialization of the numerator of* det *J*_0_ *at the chosen values. Proof*. We prove the main version first. The version for *J*_*h*_ will follow quickly from this.

Note that since *J* has at least one non-zero entry, *d*_1_ is well-defined and positive, and hence *D* is positive.

Let *W* denote numdet *J*_0_(*T*). We have that

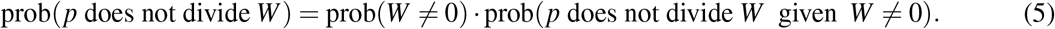

By Proposition 5.23, we have that

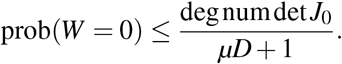

We show that the degree of the numerator of any entry of *J* is no greater than (2ν_*max*_ + 3)(*n* + *m*)*d*_1_. Fix *j* ∈ {1,…, *m*}. We first show by induction that for *k* ∈ ℕ we can write 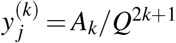,where deg *A*_*k*_ ≤ (2*k* + 1)(*n* + *m*)*d*_1_. For the base case *k* = 0, recall from Notation 5.1 we have that *y* _*j*_ = *G*_*j*_*/Q*, and deg *G*_*j*_ ≤ (*n* + *m*)*d*_1_. For the inductive hypothesis, fix *k* ≥ 0 and assume 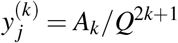 with deg *A*_*k*_ ≤ (2*k* + 1)(*n* + *m*)*d*_1_. Applying the quotient rule we have that

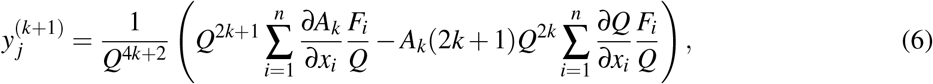

recalling from Notation 5.1 that each *f*_*i*_ = *F*_*i*_*/Q*. Since deg *F*_*i*_ and deg *Q* do not exceed (*n* + *m*)*d*_1_, we conclude the inductive step. Continuing the proof of the bound on the degrees of the numerators of the entries of *J*, note that for any *z* ∈ **x** ∪ **θ** we can write 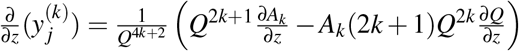 where deg *A*_*k*_ ≤ (2*k* + 1)(*n* + *m*)*d*_1_. So we can write ^∂^ (*y*^(*k*)^) = *A/Q*^2*k*+2^ where deg 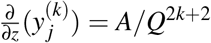.In *J*_0_ the value of *k* does not exceed ν_*max*_ so we conclude the argument.

Since *J*_0_ is square it has at most *n* + *ℓ* rows. Therefore degnumdet *J*_0_ is no greater than *D*. Thus, we have

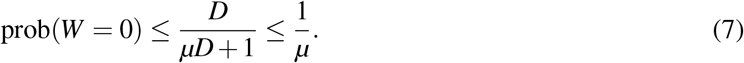

Assume that *W*≠0. Lemma 5.26 below shows that *C*≥ |*W*|. Our assumptions imply that ln*C*≥ 3 and hence *N*≥ 6. From [31, Exercise 18.18] it follows that *S* is non-empty. Using Lemma 5.24 with *M* = *W*, we have

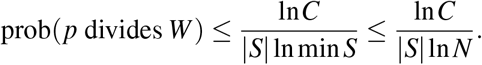

By [31, Exercise 18.18], we have |*S*| *> N/*(2 ln *N*). It follows that

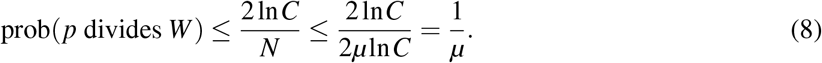

Combining (5), (7), and (8), we have our result.

We now prove the version with *J*_*h*_. Let Σ_1_ be the system obtained by adding the equation *y*_*m*+1_ = *h* to Σ and set ν_*m*+1_ = 0. Now *J*_*h*_ for Σ is equal to *J* for Σ_1_. Our result follows from the main version of the proposition.

##### Lemma 5.26.

*In the setup of Proposition 5*.*25*, ln*C* ≥ ln |*W*|.

*Proof*. For a polynomial *p* with integer coefficients, we shall define the height of *p* as ht(*p*) := max {ln(|*a*| + 1) | *a* is a coefficient of *p*}. We will use the following properties ([29, Lemma 1]): For polynomials *p*_1_, …, *p*_*s*_ in *n* + *ℓ* + *r* variables, tuple of integers *T*_0_, and partial derivative ∂, the following hold:

1. ht(∂*p*_1_) ≤ ht(*p*_1_) + lndeg *p*_1_,
2. ht(*p*_1_ + … + *p*_*s*_) ≤ max{ht(*p*_1_), …, ht(*p*_*s*_)} + ln *s*,
3. ht(*p*_1_ *p*_2_) ≤ min{deg *p*_1_, deg *p*_2_} · ln(*n* + *ℓ* + *r* + 1) + ht(*p*_1_) + ht(*p*_2_), and
4. ht(*p*_1_(*T*_0_)) ≤ max{ht(*t*_0_) | *t*_0_ ∈ *T*_0_} · deg(*p*_1_) + ht(*p*_1_).

Fix *j* ∈ {0,…, *m*} and write 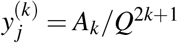,we will prove by induction that for all *k* ∈ ℕ

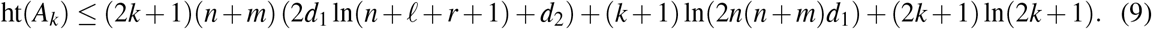

We noted earlier that deg *Q*, all deg *F*_*i*_, and deg *G*_*j*_ are no greater than (*n* + *m*)*d*_1_. It follows from this and the product height property that ht(*Q*), ht(*F*_*i*_), ht(*G*_*j*_) ≤ (*n* + *m*)(*d*_1_ ln(*n* + *ℓ* + *r* + 1) + *d*_2_). For conciseness, we will use 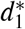 and 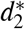 to denote these degree and height bounds, respectively.

For the base case *k* = 0, note that *A*_0_ = *G*_*j*_. For the inductive hypothesis, fix *k≥* 0 and suppose *A*_*k*_ satisfies (9). As shown in the (6), we have

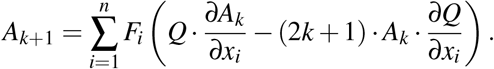

Using the derivation and product height properties, as well as the bound deg 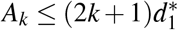.we have

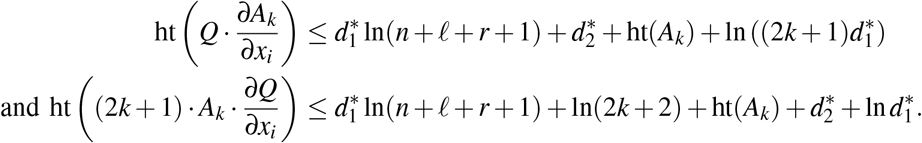

By the sum height property we have that

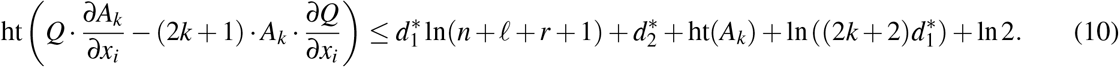

By the product height property we have

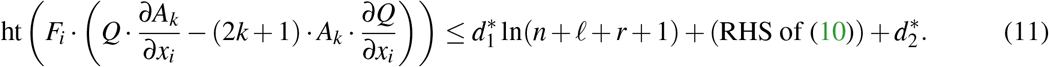

By the sum height property we have

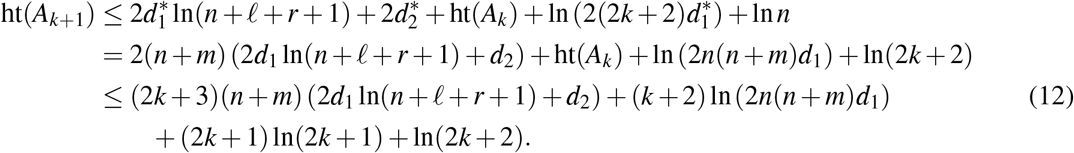

Noting that (2*k* + 1) ln(2*k* + 1) + ln(2*k* + 2) ≤ (2*k* + 3) ln(2*k* + 3), we conclude the inductive step.

Now for *z* ∈ **x** ∪ **θ**, we have that 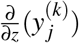 is equal to 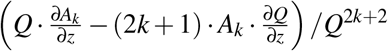. A bound on the height of the numerator is given by the right-hand side of (10). Since RHS of (10) ≤RHS of (11) ≤ RHS of (12), we see that a bound on this height is also given by the RHS of (9) with *k* replaced by *k* + 1.

Observing that the maximum value of *k* occurring for an entry in *J*_0_ is ν_*max*_ and applying the evaluation height property, we have that the numerator of each entry of *J*_0_(*T*) is bounded by

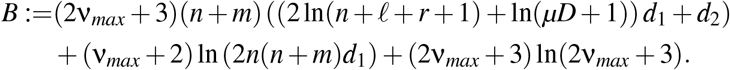

Since our bounds assume that entries in the same row have the same denominator, we have that numdet *J*_0_(*T*) = det(num *J*_0_(*T*)), where num *J*_0_ is the matrix whose elements are the numerators of *J*_0_. Henceforth we assume *J*_0_ has polynomial entries with heights bounded by *B* when evaluated at *T*. Using Hadamard’s Theorem, we have that 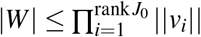, where ||*v*_*i*_|| is the square root of the sum of the squares of the elements of the *i*-th column of *J* _0_(*T*). Hence 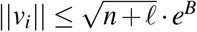.Thus

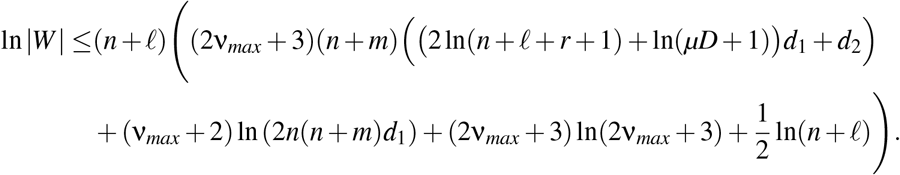

Recalling that

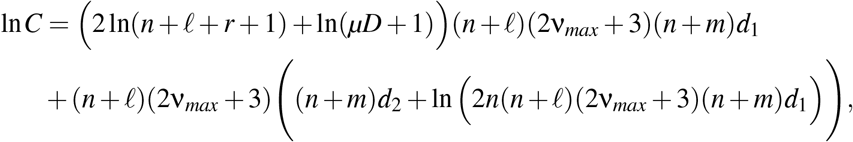

we conclude the proof.

After a set of random integers and a random prime number is chosen, we must first check that no denominator vanishes modulo the prime number before evaluating the rank of the matrix. Thus the probability that the algorithm gives the correct answer is not simply the probability that *p* does not divide numdet *J*_0_(*T*), but rather the probability that *p* does not divide numdet *J*_0_(*T*) given that *p* does not divide denomdet *J*_0_(*T*). If a random tuple *T* is chosen and used to evaluate two polynomials *A* and *B*, it is not necessarily the case that prob(*A*(*T*) ≠0) ≤ prob(*A*(*T*) ≠0 given *B*(*T*) ≠0), as the following example shows:

##### Example 5.27.

Let *k* be a positive integer and let *A*(*Z*) = *Z* and *B*(*Z*) = (*Z* − 1)(*Z* − 2) · … · (*Z* − *k*). If an integer *T* is chosen uniformly at random from {0, 1, 2, …, *k*}, then 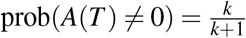 and prob(*A*(*T*) ≠ 0 given *B*(*T*) ≠0) = 0.

The following proposition shows that the conditions used for Proposition 5.25 also give a bound on the conditional probability that is relevant to our algorithms.

##### Proposition 5.28.

*Let J (resp. J*_*h*_*) be as in Algorithm 2 (resp. Algorithm 1) and suppose J (resp. J*_*h*_*) has at least one non-zero entry. Let µ, D, S, and J*_0_ *be as in Proposition 5*.*25. If a tuple of values T of the variables is chosen uniformly at random from* 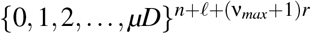 *and p is chosen uniformly at random from S, then*

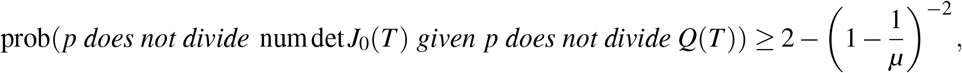

*where* numdet *J*_0_(*T*) *represents the specialization of the numerator of* det *J*_0_ *at the chosen values*.

*Proof*. Let *M* denote the sample space 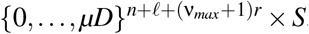. Let

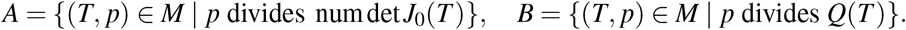

Now

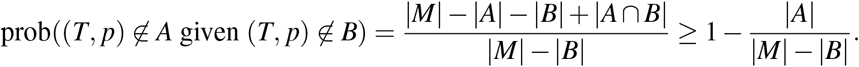

By Proposition 5.25 we have 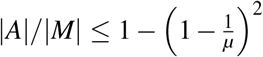.We now prove the same bound holds for |*B*|*/*|*M*|. Now deg *Q* (*n* + *m*)*d*_1_, so using Proposition 5.23 we have prob(*Q*(*T*) = 0) ≤ *D/* |*S*| = *D/*(*µD* + 1) ≤1*/µ*. By the first statement of [29, Prop. 3] with *j* = 0 we have that the height of *Q*(*T*) is no greater than (*n* + *m*)(*d*_2_ + *d*_1_(ln(*µD* + 1) + 2 ln(*n* + *ℓ* + *r* + 1))), which is no greater than ln*C*. Thus *C* ≥ |*Q*(*T*)| and if 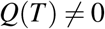 then by Lemma 5.24 we have prob(*p* divides *Q*(*T*)) ≤ ln*C/*(|*S*|*N*), which by [31, Exercise 18.18] is no greater than 1*/µ*. Thus 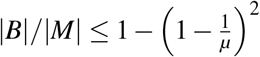.Now we have

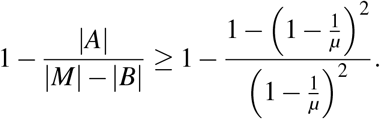

##### Proposition 5.29.

*Fix the input to Algorithm 3 (or Algorithm 4). (i) The probability that Step 1e will be reached in no more than k iterations of Step 1c is at least*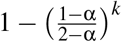.. *(ii) When Step 1e is reached the algorithm is guaranteed to terminate*.

*Proof*. (i) It was shown in the proof of Proposition 5.28 that 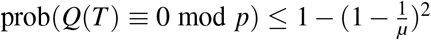,and based on the way *µ* was chosen in Step 1a it is trivial to verify that this is no greater than 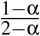.Therefore for *k* independent choices the probability that *Q*(*T*) ≢0 mod *p* for at least one of them is at least 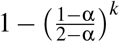.

(ii) This is obvious. □

##### Proposition 5.30.

*Fix the input to Algorithm 3. If h is* **ν***-identifiable, the probability that the output is “Yes” is at least* α. *If h is not* **ν***-identifiable, the probability that the output is “No” is at least* α.

*Proof*. Consider the following subsets of {(*T, p*) | *Q*(*T*) ≢0 mod *p*}: *Y* = {(*T, p*) | output is “Yes”}, *N* = {(*T, p*) | output is “No”}, *E* = {(*T, p*) | rank(*J*(*T*) mod *p*) = rank *J*}, and *E*_*h*_ = {(*T, p*) | rank(*J*_*h*_(*T*) mod *p*) = rank *J*_*h*_}. We have

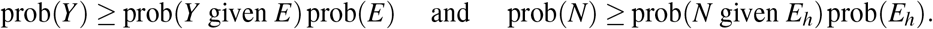

Proposition 5.28 gives us that prob(*E*) ≥ α and prob(*E*_*h*_) ≥α.

Suppose *h* is **ν**-identifiable. By Propositions 5.12 and 5.16, we have rank *J*_*h*_ = rank *J*. We show that prob(*Y* given *E*) = 1. Suppose (*T, p*) ∈ *E*. Then rank(*J*_*h*_(*T*) mod *p*) ≤ rank *J*_*h*_ = rank *J* = rank(*J*(*T*) mod *p*) ≤ rank(*J*_*h*_(*T*) mod *p*), and we deduce that rank(*J*_*h*_(*T*) mod *p*) = rank(*J*(*T*) mod *p*), so the algorithm outputs “Yes”.

Suppose *h* is not **ν**-identifiable. By Propositions 5.12 and 5.16, we have rank *J*_*h*_ *>* rank *J*. We show that prob(*N* given *E*_*h*_) = 1. Suppose (*T, p*) ∈*E*_*h*_. Then rank(*J*_*h*_(*T*) mod *p*) = rank *J*_*h*_ *>* rank *J≥* rank(*J*(*T*) mod *p*), so the algorithm outputs “No”.

##### Proposition 5.31.

*Fix the input to Algorithm 4. Let S*_*id*_ *denote the* **ν***-identifiable subset of* **x ∪θ** *and let S*_*out*_ *denote the output of Algorithm 4. The probability that S*_*out*_ = *S*_*id*_ *is at least* α.

*Proof*. Because *S*_*out*_ depends on (*T, p*) we shall use the notation *S*_*out*_(*T, p*). Consider the following subsets of{(*T, p*) | *Q*(*T*) ≢0 mod *p*}: *Y* = {(*T, p*) | *S*_*out*_(*T, p*) = *S*_*id*_} and *E* = {(*T, p*) | rank(*J*(*T*) mod *p*) = rank *J*}.

We have

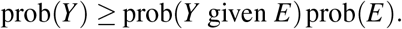

By Proposition 5.28 we have prob(*E*) ≥ α. We show prob(*Y* given *E*) = 1. Suppose (*T, p*) ∈ *E*. Let *z* ∈ **x** ∪ **θ**. Suppose *z* is **ν**-identifiable. By Propositions 5.12 and 5.16, we have rank *J*_*z*_ = rank *J*. Now rank(*J*_*z*_(*T*) mod *p*) ≤ rank *J*_*z*_ = rank *J* = rank(*J*(*T*) mod *p*) ≤ rank(*J*_*z*_(*T*) mod *p*). Since the final row of *J*_*z*_(*T*) must be (0 … 1 … 0), it follows that rank(*J*(*T*)*\z* mod *p*) = rank(*J*_*z*_(*T*) \*z* mod *p*) = rank(*J*_*z*_(*T*) mod *p*) − 1 = rank(*J*(*T*) mod *p*) − 1. Therefore *z* ∈ *S*_*out*_(*T, p*). Suppose *z* is not **ν**-identifiable. It follows from Propositions 5.12 and 5.16 that rank *J*_*z*_ − 1 = rank *J*. Because (*T, p*) ∈ *E* and the final row of *J*_*z*_(*T*) is (0 … 1 … 0) we have that rank(*J*_*z*_(*T*) mod *p*) = rank *J*_*z*_. Now rank(*J*(*T*) \*z* mod *p*) = rank(*J*_*z*_(*T*) \ *z* mod *p*) = rank(*J*_*z*_(*T*) mod *p*) − 1 = rank *J*_*z*_ − 1 = rank *J*. Therefore *z* ∉ *S*_*out*_(*T, p*). □

##### Proposition 5.32.

*Suppose that in Step 2 of Algorithm 3 (resp. Algorithm 4)* rank(*J*(*T*) mod *p*) = *n* + *ℓ. Then h is* **ν***-identifiable and the output is “Yes” (resp. every element of* **x** ∪ **θ** *is* **ν***-identifiable and the output is* **x** ∪ **θ**).

*Proof*. We first prove the statement regarding Algorithm 3. We have *n* + *ℓ* ≥ rank *J* ≥ rank(*J*(*T*) mod *p*), so rank *J* = *n* + *ℓ*. It follows that rank *J*_*h*_ = *n* + *ℓ* and by Propositions 5.12 and 5.16 we have that *h* is **ν**- identifiable. Since rank(*J*(*T*) mod *p*) rank≤ (*J*_*h*_(*T*) mod *p*) ≤*n* + *ℓ*, it follows that rank(*J*(*T*) mod *p*) = rank(*J*_*h*_(*T*) mod *p*) and the output will be “Yes”.

We now address Algorithm 4. By the preceding paragraph rank *J* = *n* + *ℓ*. Since for any *z* ∈ **x** ∪ **θ** the matrix *J\z* has only *n* + *ℓ* − 1 columns, it must be that rank *J*\*z <* rank *J* and hence by Propositions 5.12 and 5.18 each element of **x** ∪ **θ** is **ν**-identifiable. Similarly, rank(*J*(*T*) \*z* mod *p*) *<* rank(*J*(*T*) mod *p*) and hence the output is **x** ∪ **θ**. □

## Notes

### Competing Interest Statement

The authors have declared no competing interest.

### Summary of Updates

An additional analysis was conducted. Some typographical errors were corrected. A remark regarding our notion of identifiability was added.

